# Parallel genetics of regulatory sequences *in vivo*

**DOI:** 10.1101/2020.07.28.224998

**Authors:** Jonathan Froehlich, Bora Uyar, Margareta Herzog, Kathrin Theil, Petar Glažar, Altuna Akalin, Nikolaus Rajewsky

## Abstract

Understanding how regulatory sequences control gene expression is fundamental to explain how phenotypes arise in health and disease. Traditional reporter assays inform about function of individual regulatory elements, typically in isolation. However, regulatory elements must ultimately be understood by perturbing them within their genomic environment and developmental- or tissue-specific contexts. This is technically challenging; therefore, few regulatory elements have been characterized *in vivo*. Here, we used inducible Cas9 and multiplexed guide RNAs to create hundreds of mutations in enhancers/promoters and 3′ UTRs of 16 genes in *C. elegans*. To quantify the consequences of mutations on expression, we developed a targeted RNA sequencing strategy across hundreds of mutant animals. We were also able to systematically and quantitatively assign fitness cost to mutations. Finally, we identified and characterized sequence elements that strongly regulate phenotypic traits. Our approach enables highly parallelized, functional analysis of regulatory sequences *in vivo*.

## Introduction

Understanding gene regulation is fundamental for understanding development and tissue function in health and disease. Animal genomes contain diverse regulatory sequences, which are organized in contiguous stretches of genomic DNA, ranging from a few to hundreds or thousands of bases. Promoters, enhancers, and silencers act mainly on transcription while transcribed sequences (including 5′ and 3′ UTRs) regulate splicing, export, localization, degradation, and translation of mRNAs. Many regulatory sequences encode multiple regulatory functions which can cooperate, compensate and antagonize each other^1–3^. Understanding this logic requires combinatory perturbations. Moreover, a single binding site, due to fuzzy recognition motifs, may tolerate certain mutations^4–6^. The interaction of effectors with regulatory elements can additionally be modulated by sequence structure^2,5,7^, co-factors^2,3^, chemical modifications and the temporal order of binding^1,3^. Sequence activity is therefore dependent on native sequence context, cell type, tissue, development and the environment. Accordingly, phenotypic consequences of mutations in regulatory regions are difficult to predict. To understand biological functions, an approach to target regulatory sequences *in vivo*with many different mutations is required.

Although parallel interrogation of regulatory sequences has been developed in cell lines and yeast^8–12^, only a few *in vivo* approaches have been achieved in animal models. These use integration of reporters^13,14^ or injection of RNA libraries^15,16^ and therefore do not evaluate endogenous phenotypes, or are restricted to one stage of the animal life cycle. Classical genome editing by injection, now widely accessible due to CRISPR-Cas^17,18^, has enabled endogenous functional tests, but is still work-intensive and limited in scalability^19,20^.

Here we use inducible expression of Cas9 and multiplexed single guide RNAs in *C. elegans* populations to generate hundreds of targeted mutations in parallel. We targeted different regulatory regions across 16 genes and analyzed >12,000 Cas9-induced mutations to first describe characteristics of dsDNA break repair in the *C. elegans* germline and the genotype diversity that this introduces at the targeted loci. We then applied our mutagenesis approach to generate hundreds of deletions along the well-studied *lin-41* 3′ UTR, which is targeted by the miRNA *let-7*^21–24^. We developed an RNA-sequencing-based strategy to quantify the impact of each mutation on *lin-41* RNA level. Using DNA sequencing we followed the relative abundance of these different mutations over several generations to infer their phenotype. This analysis revealed a previously undescribed compensatory function between the two *let-7* miRNA binding sites. Finally, we couple the targeted mutagenesis of regulatory sequences to selection of phenotypic traits. We isolate 57 reduction-of-function alleles in 3 genes that show strong changes in phenotype, mediated by mutations in enhancer, TATA-box, and 3′ UTRs which act by both loss and gain of gene regulation.

## Results

### Inducible Cas9 to generate hundreds of mutants in parallel

To introduce many different, targeted mutations *in vivo*, we developed an approach in *C. elegans* using inducible expression of Cas9 and several multiplexed “single guide RNAs” (sgRNAs). This required only few initial injections to create transgenic animals, allowed maintenance without mutagenesis, and enabled time-controlled creation of mutated populations in parallel, with sizes only limited by culturing approaches (up to ∼10^7^ in our case). Mutant populations could then be used for various purposes. For example, they could be selected by phenotype or reporter activity, or directly analyzed by targeted sequencing to measure the impact of mutations on RNA regulation or fitness (**Fig. 1a**).

**Fig. 1.**
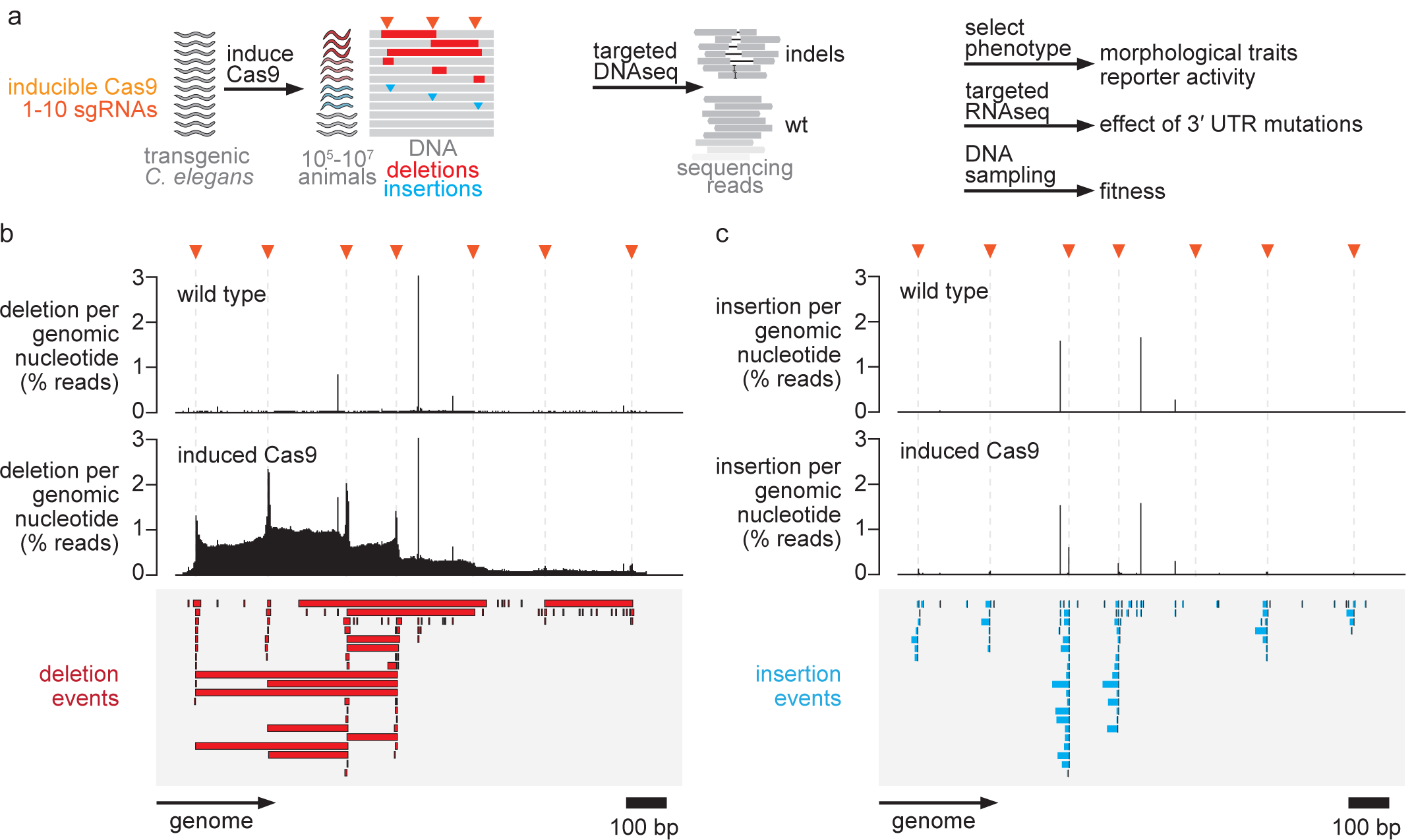
Cas9 induction to produce multiplexed *C. elegans* mutants. **a**, Outline of our approach which uses heat-shock Cas9 induction to create large “diversified” populations containing indel mutations at the targeted region. Mutated populations can be used for various downstream assays: selection according to morphological traits and reporter activity, bulk RNA sequencing to measure effects of individual 3′ UTR mutations, or DNA sampling over several generations to infer fitness of different genotypes. **b**, Example of a targeted locus (upstream of *snb-1*). The percentage of DNA sequencing reads containing deletions with respect to the total read coverage is plotted at the corresponding genomic position. Bulk worm samples were sequenced, thus 2% deletions per genomic nucleotide refers to approximately 2% of worms with a deletion at the respective nucleotide. SgRNA cut sites are indicated by orange triangles. Individual deletion events are shown below in red. **c**, Same analysis as in b) but for insertion events.

As an initial test, we generated transgenic lines with plasmids for heat shock-driven Cas9 expression and one- or multiple sgRNAs targeting a ubiquitously expressed single-copy GFP reporter. After a transient heat shock, we could observe GFP-negative animals in culture, indicating activity of Cas9. We performed a two-hour heat shock induction of Cas9 in the parents (P0) and collected progeny (F1) in a time course experiment. The highest fractions of mutants were obtained 14 – 16 hours after heat shock, with ∼50% (sg1) and ∼20% (sg2) of eggs producing GFP negative animals (**Supplementary Fig. 1a**). We obtained similar results when targeting the *dpy-10* gene and counting the characteristic Dumpy (Dpy) phenotype with the highest fractions 12 – 15 hours after heat-shock, and ∼20 – 35% of eggs producing Dpy animals (**Supplementary Fig**.**1b**).

Characteristic CRISPR-Cas9-induced mutations from 91 GFP negative animals consisted of deletions, insertions or a combination of both and originated from sgRNA cut sites (**Supplementary Fig. 1c, d**). When we used three sgRNAs within the same transgenic line targeting adjacent positions, deletions appeared around one cut site or spanned between two cut sites (**Supplementary Fig. 1d**). This indicated that pools of sgRNAs could lead to more diverse genotypes and cover more nucleotides. Most deletions induced by a single sgRNA were between 3 – 10 bp long and we observed insertion lengths between 1 – 30 bp (**Supplementary Fig. 1e**).

Homozygous animals will be produced in the F2 by heterozygous self-fertilizing F1. Additionally, since Cas9 induced in the P0 could still be active after fertilization, F1 animals could be mosaic with a wild type germline and mutant somatic cells (**Supplementary Fig. 1f**). We therefore wanted to assess how many germline mutations were generated. For this we analyzed the inheritance of GFP negative animals from F1 to F2 generations using an automated flow system and found that ∼80% of mutations were indeed germline mutations (**Supplementary Fig. 1g- i**). For the rest of our work we used such non-mosaic F2, generated by F1 germline mutations.

To analyze large mutated *C. elegans* populations in bulk, we established a targeted sequencing protocol based on long 0.5-3 kb PCR amplicons. It allowed us to sequence the complete targeted locus, to capture large deletions and to multiplex samples for sequencing (**Supplementary Fig. 2a**). This resulted in ∼200,000 – 800,000-fold read coverage at the targeted regions. We also created a software pipeline (“crispr-DART” for “CRISPR-Cas Downstream Analysis and Reporting Tool”) to handle targeted sequencing data of such amplicons and analyze the contained indels. The pipeline works with various targeted sequencing technologies and extracts and quantifies indels. The output contains html reports of coverage, mutation profiles, sgRNA efficiencies and optional comparisons between two samples. Processed genomics files from the output can then be used for more in-depth custom analyses with additionally supplied R scripts (**Supplementary Fig. 2b-d**) (“Code Availability” in **Supplementary Methods**).

To test our approach in larger scale, we induced Cas9 in 50,000 P0 animals by heat shock, and amplicon sequenced the mutated locus from bulk samples of 400,000 F2 progeny. Deletions per genomic base-pair peaked sharply around sgRNA cut sites. Pools of multiplexed sgRNA plasmids resulted in deletions spanning two or several sgRNAs (“multi cut”) in addition to smaller deletions surrounding single sgRNAs (“single cut”) (**Fig. 1b**). Insertions occurred within a few nucleotides to cut sites and were less frequent than deletions (∼1/2 – 1/10^th^) (**Fig. 1c**).

### Features of CRISPR-Cas9-induced indels and genotype diversity

To understand gene regulatory logic, ideally many different variants are produced and tested for their effects *in vivo*. For this we set out to analyze the genotype diversity produced with our approach. We targeted 16 genes at different regions with 1-9 sgRNAs per transgenic line. These genes were selected for different downstream experiments and contained one gene with a known miRNA interaction, 8 genes with strong organismal phenotypes, and 7 genes with lethal phenotypes. After Cas9 heat shock-induction we sequenced bulk genomic DNA from resultant 400,000 F2 animals with long amplicon sequencing. Together with wild type controls this produced 60 samples and data of 12,000 indel mutations for 91 sgRNAs (**Supplementary Table 1**). To measure sgRNA efficiencies, we quantified deletions +/-5bp bracketing a given sgRNA cut site. The median efficiency was ∼1.4% with most sgRNAs showing efficiencies ∼0 – 6.3% (95% CI) (**Fig. 2a**). 1.4% corresponded to approximately 5,600 mutant animals per sgRNA in our samples. We could not determine factors associated with the efficiency of sgRNAs in our system but found that lethal phenotypes were not confounding this analysis (**Supplementary Fig. 3a-d**). We then used the detected >12,000 indel mutations to characterize CRISPR-Cas9-induced dsDNA break repair outcomes in the *C. elegans* germline. We analyzed the proportion of mutation types for each sample. On average, samples contained 60% deletions, 25% insertions and 15% complex events (**Fig. 2b**). These proportions are similar for naturally occurring indels in *C. elegans*^25^ (75% deletions, 25% insertions) and human^26^ (50% deletions, 35% insertions), suggesting that natural and CRISPR-Cas9-induced dsDNA breaks undergo similar repair mechanisms.

**Fig. 2.**
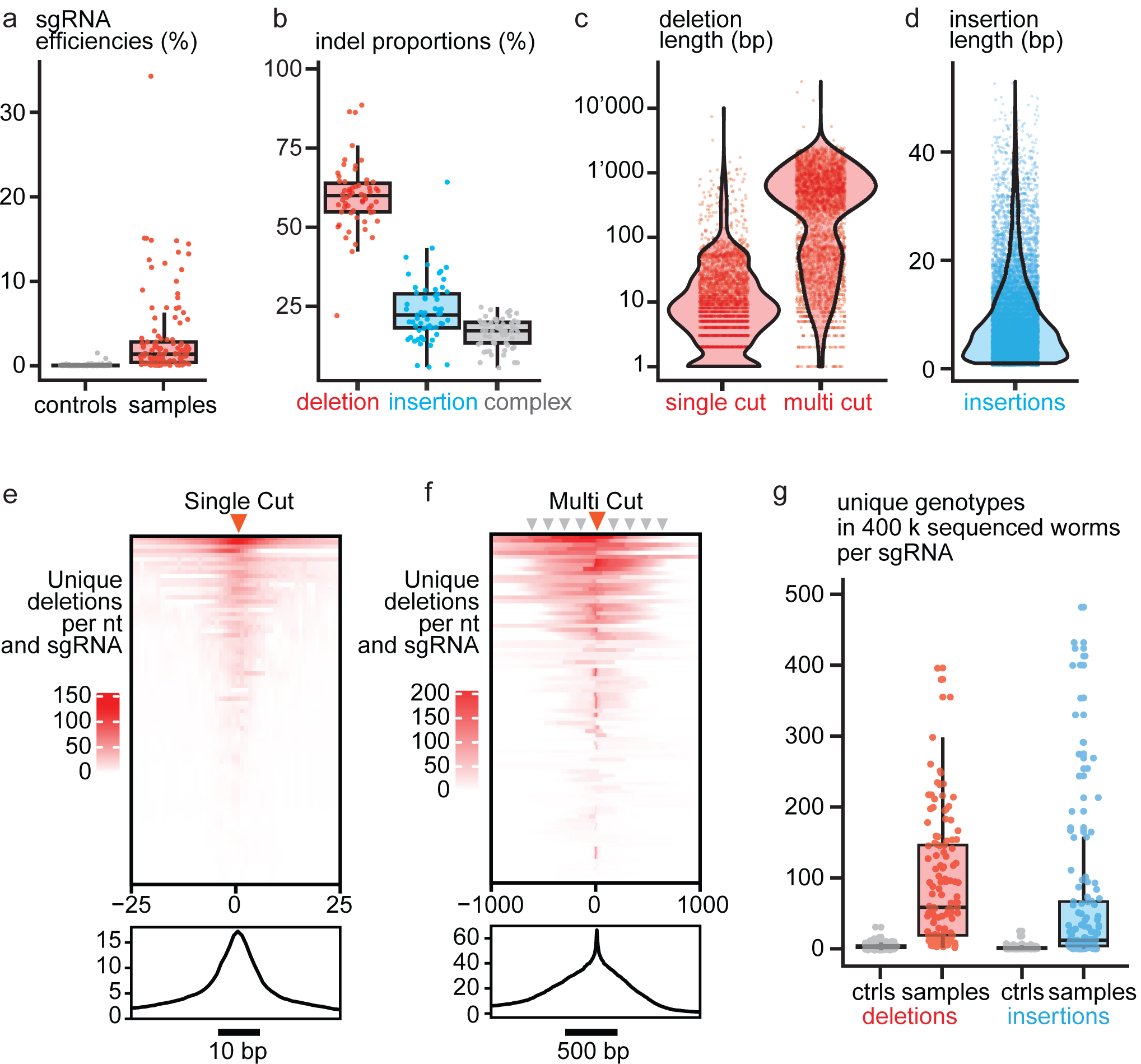
Features of sgRNA-induced mutations. Pooled data from 60 experiments (24 wild type controls, 36 samples with induced Cas9), each expressing 1-8 sgRNAs targeting one region among 16 genes (for an overview of samples see Supplementary Table 1). **a**, Efficiency measured for each sgRNA (n=127) per experiment. This is calculated by counting the fraction of reads containing deletions +/- 5 bp around the sgRNA cut site. **b**, Proportions of reads with different types of mutations detected in each experiment (n=60 experiments). Complex: reads with more than one insertion or deletion, or additional substitutions which suggest a combination of multiple events. **c**,**d**, Length distribution of mutations supported by >5 reads and >0.001% frequency, found in all experiments. Deletions were classified as multi cut deletions (n=2915) when a deletion overlapped with more than one sgRNA cut site +/- 5bp or otherwise were classified as single cut deletions (n=3169) (for a scheme describing this see Supplementary Fig. 2). Insertions supported by at least 5 reads with the exact insertion sequence at the same genomic coordinate and a frequency >0.001% (n=6616). **e**,**f**, Unique deletions (>0.001% frequency and >5 reads) per bp. Each row shows deletions per bp for one sgRNA (n=86 sgRNAs); the black curve on the bottom represents average unique deletions per nt and sgRNA. **g**, Unique genotypes detected per sgRNA in 400 k sequenced worms, classified by deletions or insertions. Each genotype is detected by >0.001% of reads mapped to the analyzed position. Ctrls n=76 cut sites, samples n=86 cut sites. Wilcoxon, p < 2.2e−16 for deletions, p < 2.2e−16 for insertions.

We computationally separated deletions into those originating from a single sgRNA (“single-cut”) or from two or more sgRNAs (“multi-cut”) (**Supplementary Fig. 3e**). The length of single-cut deletions ranged from 1 to over 100 bp, with the majority being around 5 – 25 bp. Multi-cut deletions were larger, mostly several hundred bp, as expected from the spacing between multiplexed sgRNAs (**Fig. 2c**). Most (>90%) insertions were 1 – 20 bp long although we could find insertions up to 45 bp (**Fig. 2d**). These length distributions were similar to our observations made by Sanger sequencing (**Supplementary Fig. 1e**). Insertion and deletion lengths reported from human cell lines are dramatically shorter, with the majority being 1 or 2 bp^27–30^. Longer indels in our samples can likely be explained by a higher activity of microhomology-mediated end joining (MMEJ) which uses 5 – 25 bp microhomologies and which has been reported as the main dsDNA break repair pathway in *C. elegans*^31^.

Inspection of insertions from individual genotypes revealed that most contained short sequences found in close vicinity to the insertion position (**Supplementary Fig. 4a**). Using our deep sequencing data, we systematically analyzed matches between insertions and the surrounding regions. 5-mers from insertions matched to sequences in a window+/- 13 bp around the insertion position and only in the same orientation (**Supplementary Fig. 4b-d**). Thus, our data indicate that many insertions are duplications of surrounding microhomologous sequences in the same orientation, likely the product of dissociation and re-annealing during MMEJ (**Supplementary Fig. 4e**).

Finally, we assessed the diversity of the generated mutations. We started by counting the number of unique deletions per base pair. We first studied deletions created by single-cut events for each sgRNA and found that highly active sgRNAs could generate up to 150 unique deletions (rows in **Fig. 2e**). Most deletions covered a region 10 – 12 bp surrounding the cut sites. On average, every sgRNA could generate 15 unique deletions per bp at the center of the cut site and up to an additional 5 unique deletions 5 bp away from the cut site (black line profile in **Fig. 2e**). We then studied multi-cut events. Naturally, these deletions spanned much larger regions, as defined by the spacing between two or more sgRNAs. Here we found up to 200 unique deletions per base pair and on average 20 unique deletions per sgRNA covering a region >500 bp surrounding each cut site (**Fig. 2f**). We then considered each unique indel a genotype (>0.001% frequency and >5 reads). On average, one sgRNA created 50 deletion genotypes and 10 insertion genotypes. However, some sgRNAs created up to 400 genotypes (**Fig. 2g**). Since we used several sgRNAs per transgenic line, we observed on average ∼200 insertion and deletion genotypes per sample and in efficient lines up to 1700 deletion and 1200 insertion genotypes (**Supplementary Fig. 5a**). More efficient sgRNAs resulted in a higher number of unique deletions (**Supplementary Fig. 5b**). Transgenic lines expressing more sgRNAs showed more unique deletion genotypes, possibly because of an increased chance of containing efficient sgRNAs and the combined activity of multiple sgRNAs (**Supplementary Fig. 5c**). Together these data show that inducible expression of Cas9 with multiplexed sgRNAs can induce a variety of mutations to study hundreds of regulatory variants in parallel. This includes small deletions to target individual elements at nucleotide resolution, large deletions to interrogate combinatory interactions and insertions to change spacing between sites or duplicate existing sequences.

### Quantifying mutant gene expression and fitness

A major challenge to understand gene regulation is the interaction of different elements. Especially in 3′ UTRs, which can act on all levels of gene expression, this can be difficult. To simultaneously measure mRNA levels for all generated 3′ UTR deletions within large *C. elegans* populations, we developed a targeted RNA sequencing strategy. As a proof of principle, we tested it on a microRNA-regulated mRNA. The *lin-41* mRNA is regulated by *let-7* microRNAs which bind two complementary sites in the 1.1 kb long 3′ UTR (LCS1 and LCS2, 22 and 20 nucleotides long, separated by a 27 nt spacer)^21,22,24,32,33^ (**Supplementary Fig. 6a**). Although studies with reporter plasmids showed that each binding site could not function on its own^22^, other studies concluded that each site could recapitulate wild-type regulation when present in three copies^23^. Prior to our work, compensation had not been tested for each site, in the native sequence context, or at natural expression levels. Therefore, we targeted the *lin-41* 3′ UTR with a pool of 8 sgRNAs and two sgRNA pairs close to the LCSs (**Fig. 3a**). *Lin-41* down -regulation occurs with *let-7* expression in the developmental stages L3-L4^21,32–34^. To measure *let-7*-dependent regulation, we collected RNA from bulk worms at L1 and L4 stages. We extracted L4 stage RNA after complete *lin-41* mRNA downregulation by *let-7*^35^ and before the occurrence of the lethal vulva bursting phenotype^24^ (**Supplementary Fig. 6b**) (see **Methods**). We then sequenced *lin-41*-specific cDNA with long reads covering the complete 3′ UTR (**Supplementary Fig. 6c**). Each read contained full information on any deletion present in the RNA molecule, while the number of reads supporting each deletion could be used to estimate RNA expression level. To determine *let-7*-dependent effects, we then analyzed how different deletions affect RNA abundance at L4-relative to L1 stage.

**Fig. 3.**
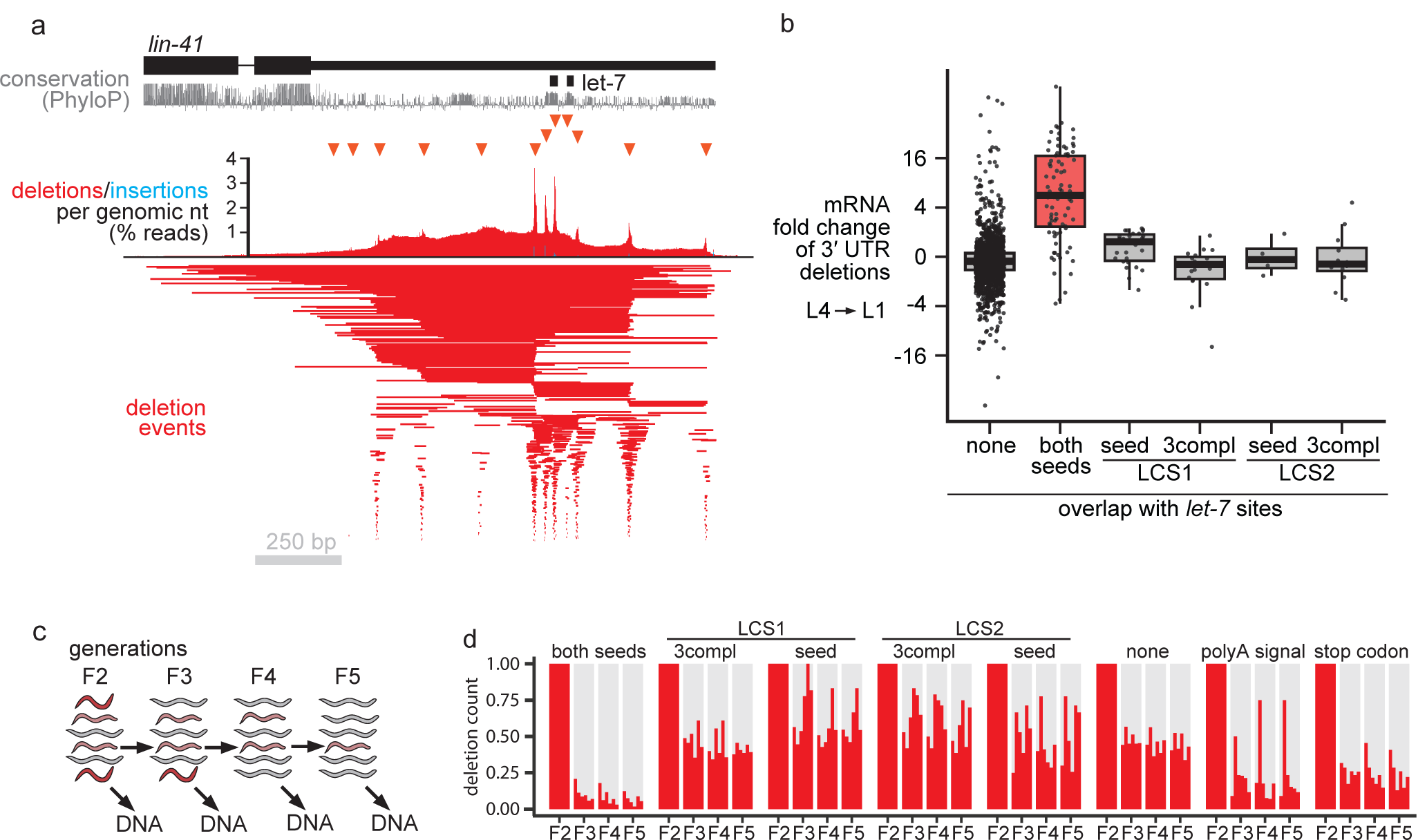
Analysis of the *lin-41* 3′ UTR shows redundancy of the two *let-7* miRNA binding sites for RNA regulation and fitness. **a**, The *lin-41* 3′ UTR locus after targeted mutagenesis with three different lines (sg pool, sg1516, sg2627). Deletions of three lines were pooled and analyzed together. Deletion events (n>900) supported by >0.001% reads. **b**, Relative fold change of deletions detected in *lin-41* 3′ UTR full-length cDNA between L1 and L4 developmental stages. We considered only deletions that are detected in both L1 and L4 RNA samples. **c**, Generations were separated by sedimentation and worms were sampled for DNA extraction. **d**, Reads supporting deletions in bulk genomic DNA of consecutive generations, relative to the first (F2) generation. At each generation shown are all six samples (sg pool, sg1516, sg2627 grown at 16 and 24°C)

We observed an average 4-fold up-regulation of *lin-41* at larvae stage L4 only when both *let-7* seed sites were mutated by deletions (**Fig. 3b**). No significant effect was observed from deletions affecting either LCS1 or LCS2 seeds alone. The 4-fold regulatory effect is consistent with the known magnitude of down-regulation in the natural context^22,32,33^ or the up-regulation when disrupting both *let-7* interactions^24,36,37^ (2 – 4-fold). A very slight and non-significant up-regulation was observed for LCS1. Interestingly, LCS1 has a longer seed pairing (8 nt) than LCS2 (6 nt and a G-U pair) and was detected with more reads in an *in vivo* miRNA proximity ligation approach^38^. In addition 3xLCS1 acted stronger than 3xLCS2 in a reporter assay^23^. As an independent approach, we used UMAP on the k-mer content of long cDNA reads to obtain clusters of genotypes^39^. These data also suggest that RNA molecules transcribed from clusters with large deletions overlapping both LCS seeds were detected with more reads in L4 stage compared to L1 stage animals (**Supplementary Fig. 6d-f**).

To assign fitness to individual mutations in a controlled environment, we established measurements on genotype abundance over several generations. For this, we sampled genomic DNA of consecutive generations (**Fig. 3c**). Disrupting *let-7* regulation of *lin-41* is known to result in lethal developmental defects^21,24,32,37,40^. Deletions in the *lin-41* 3′ UTR which overlapped with both LCS seeds quickly disappeared from the population and were mostly absent already after one generation (**Fig. 3d**). Consistent with our observed absence of an effect on RNA expression, deletions affecting either one of the two LCS alone were not significantly depleted. Our results indicate that the two *let-7* complementary sites can largely compensate each other’s loss under laboratory conditions. We conclude that parallel mutagenesis coupled with targeted RNA or DNA sequencing can be used to analyze function and interactions of regulatory elements *in vivo* directly from large populations in bulk.

### Gene regulatory sequences which affect phenotypic traits

Next, we w anted to isolate regulatory sequence variants which affected animal phenotype. This could be useful to discover functional elements, provide starting points to study mechanisms, and to obtain animals with desired phenotypic traits. Such an approach would also capture any functional sequences regardless of the level, time or tissue of regulation, which can be hard to predict. We proceeded to select worms based on strong organismal phenotypes affecting animal movement and body shape (Unc, Rol, Dpy) and to identify the causative mutations. We targeted a predicted enhancer^41^, three promoters and all 3′ UTRs of 8 genes with known loss-of-function phenotypes and screened 35,000 – 45,000 animals for each of these regions (**Supplementary Table 2**). To determine which mutations were initially present in the screened population, we performed targeted sequencing on siblings (**Supplementary Fig. 8a, b**). Initially, we isolated several mutants with large deletions (>500 bp) that disrupted the coding frame or the polyadenylation signal (AATAAA) (**Supplementary Fig. 8c, d**). Similar large-scale, on-target deletions have previously been described in cell lines^42,43^ and mice^44^. We also found large insertions (<250 bp) which originated from within +/-1 kb of the targeted region, or from loci on other chromosomes (**Supplementary Fig. 8c, d**). We found such large deletions or insertions in 5 out of 8 screened genes, demonstrating that for these genes our screen was sensitive enough to detect animals with affected phenotypes (**Supplementary Table 2**).

From the screen we isolated 57 reduction-of-function alleles in 3 genes (*egl-30, sqt-2, sqt-3*) and none from the other 5 genes (*dpy-2, dpy-10, rol-6, unc-26, unc-54*) (**Supplementary Table 2**). Deletions, insertions and complex mutations were represented equally among isolates (**Supplementary Fig. 8e**). The observed phenotypic traits showed complete penetrance and we scored their expressivity which differed between mutations. We found that several mutations in the 3′ UTR of *egl-30* lead to Uncoordinated (Unc) phenotypes. In 7/11 mutants, a region circa 100 bp downstream of the STOP codon was affected. The smallest deletion was 6 bp (**Fig. 4a**) (**Supplementary Fig. 8g**). We also found mutations with Roller (Rol) phenotypes overlapping a putative *sqt-2* enhancer predicted from chromatin accessibility profiling^41^ (**Supplementary Fig. 8f**). We also targeted *sqt-3*, a gene associated with three distinct morphological traits^45,46^ (Dpy, Rol and Lon). In total we isolated 39 alleles. 13 mutations upstream of *sqt-3* likely affected transcriptional initiation, with 11/13 overlapping the predicted TATA-box (**Fig. 4b**). In line with the Rol phenotype, which indicates reduction-of-function, pre-mRNA and mRNA levels were both reduced to ∼50% (**Supplementary Fig. 9d**). The remaining *sqt-3* alleles were 3′ UTR mutations. Almost all (25/26) were insertions or complex mutations originating at sg2 (**Fig. 4c**). The only deletion overlapped with the polyadenylation signal (AAUAAA). We knew from sequencing siblings that sg2 was very efficient (∼25%) and that in our samples various deletions covering the 3′ UTR were present. We therefore isolated 13 distinct non-Rol mutants using direct PCR screening (**Supplementary Fig. 9a, b**). Despite containing deletions or insertions originating at the efficient sg2, these animals showed the wild type non-Rol trait (**Fig. 4d**). We did follow-up experiments with one of the 25 insertion alleles, *sqt-3(ins)* and determined that mRNA levels were reduced post-transcriptionally to ∼50% (**Supplementary Fig. 9c, d**). Since deletions in this region were well tolerated (non-Rol), we concluded that the isolated Rol mutations likely resulted from a gain of repressive sequence which led to the observed reduction of mRNA. Interestingly the polyA mutant *sqt-3(polyA)*, for which mRNA levels were equally reduced to 50%, showed a much weaker Rol phenotype, with only slight bending of the head (**Fig. 4c, Supplementary Fig 9d, e**). This suggests that additional mechanisms besides mRNA down-regulation might reduce protein output in *sqt-3(ins)*.

**Fig. 4.**
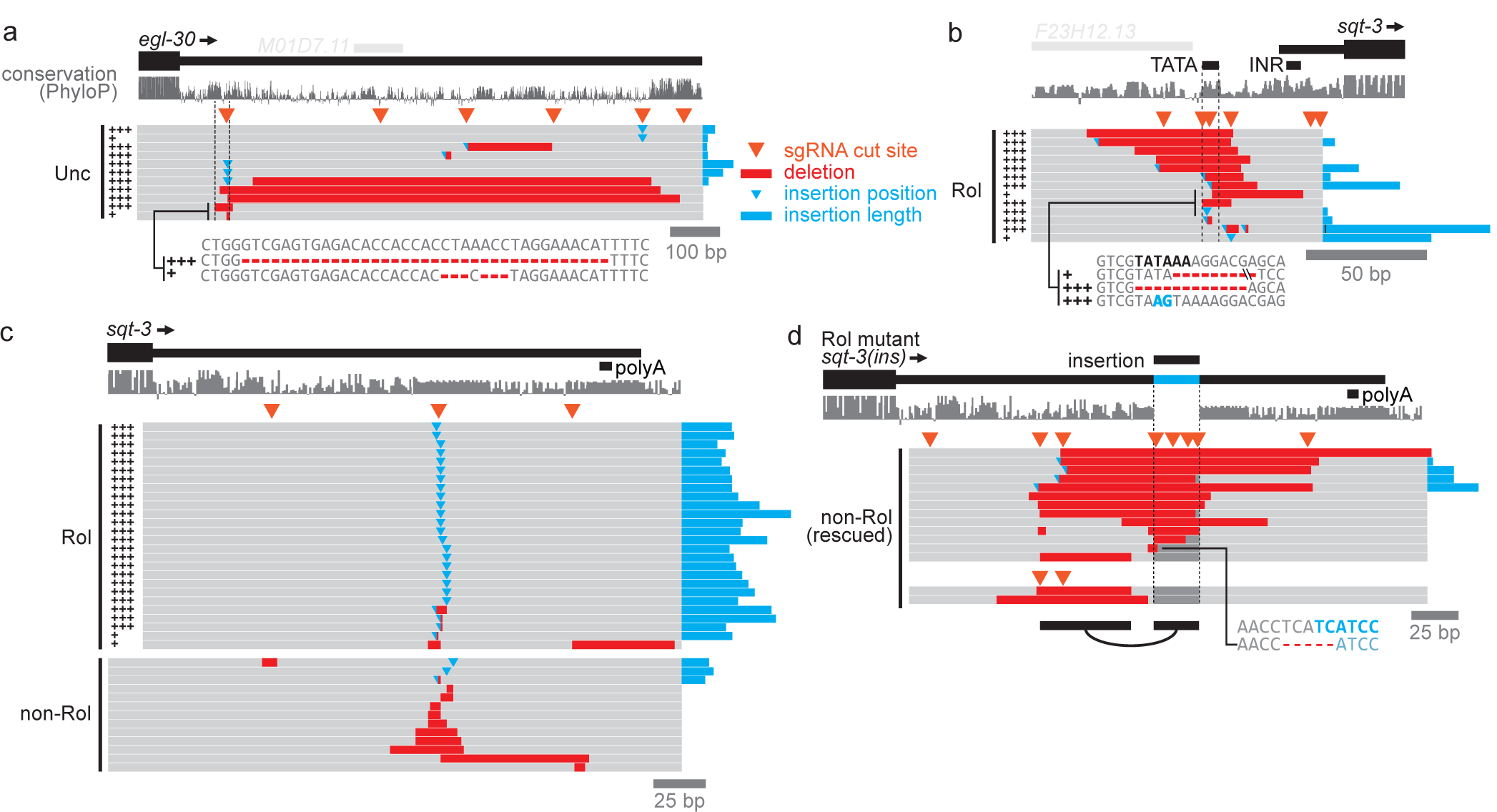
A screen finds regulatory mutations strongly affecting phenotype. Shown are genotypes of strains which were isolated according to phenotypic traits after targeting regulatory regions. Phenotypes showed complete penetrance (n>300 animals) and expressivity was scored as indicated by +, ++, or +++ (n>300 animals). **a**, Eleven mutations along the *egl-30* 3′ UTR which show slight or strong Uncoordinated (Unc) phenotypes. **b**, Thirteen mutations upstream of *sqt-3* which show a Roller (Rol) phenotype. Most (11/13) overlap with the predicted TATA-box. **c**, Mutations in the *sqt-3* 3′ UTR which show a Rol phenotype or which are tolerated (non-Rol). Almost all (25/26) Rol mutants contained insertions. **d**, Fifteen mutations, mostly deletions, which rescue the Rol phenotype of one insertion allele *sqt-3(ins)*. While most overlapped with the insertion isolated for a Rol phenotype, three deletions affected a region upstream (black bars on the bottom indicate the uncovered compensatory interaction).

To define the repressive sequence elements, we targeted the inserted sequence with several sgRNAs and screened for non-Rol “revertants” in which the wild type trait was restored. 12/13 revertants contained deletions overlapping with the insertion, with the smallest being 5 bp (**Fig. 4d**). A restored wild type trait likely resulted from restored expression levels and indeed mRNA levels in two independent revertants were restored to normal (**Supplementary Fig. 10b**). Overall, predicted RNA secondary structures did not change, suggesting other factors cause the Rol phenotype of *sqt-3(ins)* (**Supplementary Fig. 10c**). Finally, we found, using sequence transplantations into hypodermally expressed *dpy-10* and neuronally expressed *unc-22*, that repression was dependent on 3′ UTR context and observable also in neuronal tissue (**Supplementary Fig. 10d**). This implies that we isolated a general repressive/destabilizing RNA element.To discover interacting 3′ UTR elements we had included sgRNAs for the remaining 3′ UTR. This revealed a compensatory deletion upstream of the insertion, which was able to revert the Rol phenotype. We isolated two more additional alleles after using sgRNAs specific for this region (**Fig. 4d**) (**Supplementary Fig. 10a**). Surprisingly, mRNA levels were not restored (**Supplementary Fig. 10b**). This points to an alternative mechanism of restored protein function, for example on translational level, or affecting mRNA at a different developmental time point.

Overall, these results demonstrate that parallel genetics and selection by phenotype can be used to obtain specific phenotypic traits, to find functional sequences, and to discover unexpected regulatory interactions *in vivo*.

## Discussion

In this study we develop a general approach for parallel genetics of regulatory sequences in vivo, using inducible expression of a CRISPR-nuclease and multiplexed sgRNAs. Large “diversified” populations can then be used for comprehensive analysis using direct deep sequencing or for selection by phenotypic traits. This allows directly linking regulatory genotypes with phenotypes. We demonstrate this in the model organism *C. elegans* but believe it is similarly applicable in other animals or plants which allow transgenesis and inducible expression of genome editors.

As we show, sgRNA efficiencies around 1.5 % are sufficient to analyze effects of mutations on gene regulation and phenotype when coupled with deep sequencing and manual- or automated selection of animals from large populations. However, higher efficiency would be desired for improved comprehensive testing. This could be achieved with more advanced expression systems^47^ and optimization of sgRNA promoters for high germline expression. Alternative induction systems (e.g. Auxin, FLP/FRT, Cre/lox, Gal/UAS) could enable continuous and germline-specific Cas9 expression to increase efficiency and allow directed evolution experiments. Our targeted sequencing protocol can capture long deletions, uses the same amplicon for the whole locus, and allows sample multiplexing. Unique molecule counting methods^48^ for long reads^49,50^ could be incorporated to reduce PCR biases. Established protocols^28^ are available for shorter (100-300 bp) target regions. We assumed that each animal in bulk samples contributed equally to the extracted genomic DNA. In the future, barcoding to determine genotypes of individual animals could be developed with plate-based or split-pool^51,52^ methods. Our method only works at nucleotide resolution close to the sgRNA cut sites. To allow denser tiling of regions with mutations, CRISPR-nucleases with alternative or dispensable PAM requirements could be used^17,53,54^. Although indels are applicable to regulatory regions and even coding sequences^55–57^, point mutagenesis would enable fine mapping of regulatory nucleotides and amino acids. This exciting possibility could be opened up by implementing hyperactive base^58–61^- or prime editors^62–64^. Alternatives to CRISPR-Cas could be developed based on inducible expression of cassette exchange^65,66^ or targeted transposons^67,68^. Targeting several independent loci in one step might be applied to candidate regulatory elements (e.g. miRNA targets, enhancers), screening candidate genes (e.g. from networks, pathways^69^, other assays^70^), and for synthetic co-evolution of several loci^71^.

Indel data from high throughput genome editing in human cell lines led to insights into dsDNA break repair outcomes^27–30,72–75^. Whether these data align with outcomes *in vivo* or in the germline has not been well studied. We found longer indels and less 1nt templated insertions in our data. This can be explained with higher activity of microhomology-mediated end joining (MMEJ), which has been shown to be the main dsDNA break repair pathway in *C. elegans*^31^. Mutations typical for MMEJ have been implicated in diseases^76^ and microhomologies allow predicting the outcomes of CRISPR-Cas9 editing^27–29,72–75^. Deep mutation profiles in *C. elegans* could be used to study MMEJ *in vivo*.

In the evolution and determination of phenotypes gain or loss of regulatory sequences is important^1,77,78^ and can be modeled as a gradual process by single nucleotide changes^6,77–81^. Although indels occur less often than single nucleotide mutations naturally, their impact can be more severe. Deletions have already been found to destroy or generate regulatory elements during evolution^77,78^. Insertions templated from regions around dsDNA breaks during DNA repair could duplicate functional sequence elements and might be an underestimated force in the evolution of regulatory sequences or disease mutations.

In line with previous observations we found that the two *let-7* complementary sites repress *lin-41* ∼4-fold^22,24,32,33,36,37^. Furthermore, we uncovered a previously undescribed compensatory interaction. Previous studies concluded either that one LCS could not recapitulate wild-type repression at all^22^ or tested only multiple copies of each site^23,82^. These studies were done with reporter plasmids and therefore not in the native sequence context (which affects target site accessibility^23^), or at natural expression levels (which affects miRNA-target ratios^37^). We found a slight but non-significant effect on RNA regulation when disrupting LCS1. Together with previous studies^23,38^ this indicates that LCS1 could act more strongly than LCS2, and that both sites cannot completely compensate each other’s loss. Since our method might lack the sensitivity to detect more subtle effects from deletion of each LCS alone, future studies would be needed to test this thoroughly.

Our screen for sequences that affect phenotypic traits doubles the phenotypic regulatory alleles registered in the last forty years at Wormbase. Due to the mutagenesis efficiency and because we screened for strong phenotypes, we did not saturate and likely missed many mutations. We expect that higher efficiencies would likely lead to mutations affecting expression and phenotype also for the genes for which we did not isolate any mutations. To determine comprehensively which mutations are tolerated by a locus, even higher efficiencies would be needed. Some mutations were likely affecting known regulatory sequences (*sqt-3* TATA-box), while for others (*sqt-2* enh, *sqt-3* and *egl-30* 3′ UTR) we did not determine a mechanism. However, such mutants can be a starting point for future studies on regulatory mechanisms using biochemical, computational and genetic approaches. Additionally, the approach described here can be applied to isolate alleles with altered function (reduction-of-function, gain-of-function) and to obtain animals with desired phenotypic traits. Furthermore we discovered compensatory deletions in the *sqt-3(ins)* 3′ UTR that restored the wild type phenotype. Although not restoring mRNA levels, and likely acting by an alternative mechanism, we uncovered these by screening for the animal phenotype. Our results highlight the possibility to uncover unexpected regulatory interactions using *in vivo* interrogation and readout of phenotype. Such interactions should be detectable, even if they act across different developmental time points, tissues or cell types.

Altogether we believe the approach presented here will help in understanding how gene regulatory logic affects phenotype *in vivo*.

## Supporting information

Supplementary Materials (oligos, plasmids, sgRNAs, injection mixes, amplicons, strains)

## Methods

### Maintenance of animals

C. elegans were maintained on NGM plates with Escherichia coli OP50 as originally described^83^ at 16, 20 or 24 °C. Plates for hygromycin resistant transgenic animals were modified by adding working stock solution of 5 mg/mL Hygromycin B (Thermo Fisher Scientific, cat. 10687010) in water onto plates before use, to a final concentration of 75 μg/mL NGM. For standard 6 cm plates with 10 mL NGM that would be 150 μL of 5 mg/mL Hygromycin working stock solution.

### Strains

A complete list of strains can be found in a supplementary table. The wild-type strain N2 Bristol^84^ was used to create transgenic lines for experiments. In a screen for phenotypes we isolated several mutants and revertants for different regulatory regions. For initial tests we generated a *his-72* c-terminal GFP knock-in strain (NIK123) which we crossed into a strain expressing *Peft-3:tdTomato:H2B* from a single copy insertion (EG7927^68^) resulting in a GFP/RFP expressing strain (NIK124) for automated quantifications and sorting using the Copas Biosorter.

### Plasmid construction

A list of all plasmids created or used in this study can be found in a supplementary table. The plasmid for heat-shock inducible *Streptococcus pyogenes* Cas9 expression (pJJF152) was created by Gibson assembly^85^ of a previously published *C. elegans* optimized SpCas9^86^ (“Friedland Cas9”), with the *hsp-16*.*48* heat-shock promoter and the *unc-54* 3′ UTR.

Plasmids for sgRNA expression were cloned as previously described using one of two published backbones (pMB70^87^ or pJJR50^88^). For this, 5-10 μg of backbone was digested using 1 μL Fastdigest Eco31I (aka BsaI, Thermo Fisher Scientific, cat. FD0293) or Fastdigest BpiI (aka Thermo Fisher Scientific, cat. FD1014) at 37°C for 2-6 hrs, separated from undigested plasmid on a 1.5% Agarose/TAE gel, and extracted using the Zymoclean Gel DNA Recovery Kit (Zymo Research, cat. D4002), according to the instruction manual. Two complementary DNA oligonucleotides containing the spacer sequence, plus an optional 5’ G for optimal U6 promoter expression, and 4 nucleotide overhangs for ligation into the backbone were phosphorylated and annealed in a thermocycler. This reaction contained 1 μL of each oligo (at 100 μM), 1 μL of 10x T4 DNA ligase buffer (Thermo Fisher Scientific, cat. EL0011), 1 μL T4 PNK (Thermo Fisher Scientific, cat. EK0031) and 6 μL water and was incubated 37°C 30 min, 95°C 5 minutes and cooled down at -0.1 °C/second to 25°C. Sample was diluted 1:200 in water and 1 μL was used for ligation with 70-130 ng of linearized backbone, 1 μL of 10x T4 DNA ligase buffer and 1 μL of T4 DNA ligase (Thermo Fisher Scientific, cat. EL0011) and water to a volume of 10 μL. Ligation was performed at room temperature for 1 hr or overnight. 5 μL were then transformed.

The HDR repair template plasmid used for the his-72::GFP knock-in was prepared as described previously^89^.

For transformation and amplification, we used DH5alpha Mix & Go Competent Cells (Zymo Research, cat. T3007) in all the above clonings except for the *his-72::GFP* repair template which required ccdB resistant bacteria for which we used One Shot ccdB Survival (Thermo Fisher Scientific, cat. A10460). DNA extractions by miniprep were done with the ZymoPURE Plasmid Miniprep kit and elution with water (Zymo Research, cat. D4208T).

### sgRNA design

sgRNAs were designed manually using *C. elegans* genome version ce11 and the plasmid editor Ape (A plasmid Editor, M.W. Davis, https://jorgensen.biology.utah.edu/wayned/ape/) and evaluated using the E-CRISP web application (http://www.e-crisp.org/E-CRISP)^90^. For regulatory regions of interest, we aimed at a regular spacing between target sites, dense coverage and as little as possible predicted off-targets with less than three mismatches. A detailed list of sgRNA sequences, together with their characteristics, efficiency prediction scores and predicted off-targets, can be found in a supplementary table.

### Generation of transgenic *C. elegans*

Extra-chromosomal array transgenes were generated by standard procedure using micro-injection into the gonad^91^. A detailed list of injection mixes and their composition can be found in a supplementary table. The injection mix usually contained plasmids for heat-shock inducible Cas9 (pMB67^87^ or pJJF152) at 50 ng/μL, 1-10 sgRNAs (using the backbone pMB70^87^ or pJJR50^88^) at 10-50 ng/μL, a visual co-injection marker expressing mCherry in the pharynx (pCFJ90^92^) at 5 ng/μL, and hygromycin resistance (IR98^93^) at 3 ng/μL. Independent lines were created from F1 animals selected for pharynx expression of the mCherry co-injection marker. Lines were maintained on Hygromycin selection plates as described above.

### C-terminal GFP knock-in of *his-72*

C-terminal GFP knock-in of *his-72* was performed as described previously using a self-excising selection cassette^89^.

### Biosorter

Automated measurement of GFP negative animals in F1 and their F2 progeny. *His-72::GFP* was targeted with sg1, sg2, pool1 (sg2, 3, 4, 6, 8) or pool2 (sg3, 5, 8). F1 generation was collected by bleaching 12 hrs after heat-shock. These were either measured on the Biosorter flow system at larvae stage L3 or grown to adulthood to collect F2 generation which was then also measured at larvae stage L3. The number of analyzed worms per sample was between 1’662 and 21’983 worms.

### Small-scale mutagenesis by Cas9 heat shock induction and time course

20-40 egg-laying adults were transferred to small 6cm NGM plates with OP50 *Escherichia coli* and without Hygromycin. Plates were placed in a programmable incubator “Innova 42” (New Brunswick Scientific/ Eppendorf) at 20°C. Heat shock was applied for 2 hours at 34°C, followed by 20°C. For time course experiments adults were transferred to new plates using a picking tool at regular time intervals (14, 16, 18, 20, 22, 43 or 12, 15, 18, 21, 48 hrs) after heat shock to analyze eggs laid within each interval.

### Developmental synchronization

Synchronized L1s were obtained by bleaching, as previously described^94^. Egg-laying animals were washed off plates in 50 mL M9 buffer (42 mM Na_2_HPO_4_, 22 mM KH_2_PO_4_, 86 mM NaCl, 1 mM MgSO_4_) and settled for 10 minutes. M9 was aspirated until a remaining volume of 7.5 mL. Then 1 mL 12% NaClO and 1 mL 5 M NaOH were added. Worms were incubated under gentle rotation, vortexed briefly after 4 minutes and incubated under constant observation for another 3 minutes. Bleaching was stopped by addition of 40 mL M9 when circa 50% of animals were dissolved. Eggs were then pelleted by centrifugation at 1200 g for 1.5 minutes and washed two more times using M9, centrifugation and decanting. Finally, eggs were resuspended in circa 4 mL M9 and left shaking at 16 °C overnight for at least 12 hours to allow hatching and developmental arrest of L1 larvae. Larvae concentration was then counted in triplicates and the desired amount was dispensed on plates with food to begin synchronized development.

### Large-scale mutagenesis by Cas9 heat shock induction

Before the experiments, animals were maintained 5-25 generations in culture under Hygromycin selection to ensure expression of transgenes. Expression was indicated by Hygromycin resistance and the visual mCherry co-injection marker expressed in the pharynx. For all experiments three independent lines from the same injection mix were used. For transient heat shock induction of Cas9, synchronized populations were seeded on large 15 cm NGM plates with food and without Hygromycin. Plates with egg-laying adults (P0) were placed in a programmable incubator “Innova 42” (New Brunswick Scientific/Eppendorf) at 20°C and 34°C heat shock was applied for 2 hours. Plates were kept at 20°C for 12 hrs and eggs were collected by bleaching as described above for developmental synchronization. Hatched larvae, arrested at the L1-stage, the first generation after Cas9 induction (F1), were then again seeded on large NGM plates with food for synchronized development until egg-laying, to collect the next generation (F2) by bleaching. We used this F2 generation for all experiments to ensure non-mosaic animals generated by F1 germline mutations. We seeded 50,000 P0 for Cas9 induction at 24°C on Hygromycin (25,000 / big plate), and 100,000 F1 at 16°C (25,000 / big plate). 400,000 F2 were frozen for genomic DNA extraction to determine introduced indel mutations. The remaining F2 were used for experiments described below.

### Genomic DNA extraction

Genomic DNA was obtained using worm lysis, phenol-chlorophorm extraction and ethanol precipitation. Worms were washed once in 50 mL M9 buffer and frozen in 1 mL M9. After thawing, M9 was removed and 100 μL of TENSK buffer (50mM Tris pH 7.5, 10 mM EDTA, 100 mM NaCl, 0.5% SDS. 0.1 mg/mL proteinase K, 0.5% ß-Mercaptoethanol) was added. Sample was incubated for 1.5 hrs at 60 °C while shaking at 1000 rpm on a benchtop heating block. 300 μL of water was added, followed by 400 μL phenol/chlorophorm/isoamylacohol pH 8.0 (Carl Roth, cat. A156). Sample was mixed by shaking the tube and centrifuged for 10 min. at 15’000 g at room temperature. The upper aqueous phase, circa 350 μL, was transferred to a new tube and an equal volume of chlorophorm was added. After additional centrifugation 10 min. at 15’000 g at 4°C, the upper aqueous phase was transferred to a new tube, and 2 μL glyco blue added. This was followed by addition of 30 μL 3M NaAc (pH 5.2-6) and 1 mL pure ethanol. Samples were centrifuged for 10 min. at full speed and 4°C in benchtop centrifuge. Pellet was washed once with 70% ethanol and resuspended in 25 μL water at 50°C for 30 min. Then 0.25 μL RNAse I (10 U/μL, Thermo Fisher Scientific, cat. EN0601) was added and incubated for 30 min. at 37°C. DNA concentration was determined on a Nanodrop ND-1000 (Thermo Fisher Scientific) and diluted to 50-200 ng/μL in water.

### DNA large amplicon sequencing

Amplicons were designed so that they contained all the regions of a gene targeted in our experiments. 0.5 – 3 kb amplicons were large enough that deletions between the outermost sgRNAs would not change the amplicon size by more than 10% to avoid more efficient amplification of templates with large deletions. Furthermore, large amplicons should capture the reported large deletions missed by 100-300 bp amplicons of other workflows. Primers used for amplification together with annealing temperature and resulting amplicon sizes can be found in a supplementary table.

Genomic DNA concentration was fluorimetrically quantified using Qubit dsDNA HS kit (Thermo Fisher Scientific, cat. Q32854). For PCR reactions we used 100 ng template DNA. We calculated that 100 ng of genomic DNA equals >90 million *C. elegans* genomes and therefore represented all animals (maximal 2 million) in a sample.

50 μL PCR the reactions were set up as follows. Phusion HF polymerase (New England Biolabs, cat. M0530L) 0.2 μL, 5X HF buffer 10 μL, dNTP mix 1 μL, forward and reverse oligos at 10 μM 5 μL, water 32 μL, and template DNA. Samples were incubated at 98°C 3 min, followed by 35 cycles of 98°C 15 sec, 58-72 °C 30 sec, 72 °C for 7 min with a final elongation at 72 °C for 7 min. PCR reactions were analyzed on agarose gels to ensure successful amplification.

Cleanup was then done by either agarose gel or SPRI beads. For gel-based cleanup 1.5 % Agarose/TAE gels were run and bands were excised with circa +/-500 bp, to also include products with deletions or insertions. DNA was recovered from agarose gel using the Zymoclean Gel DNA Recovery Kit (Zymo Research, cat. D4002). For SPRI beads cleanup and no size selection we used AMPure XP Reagent (Beckman Coulter, cat. A63881). 0.8 x volume of beads were added to PCR reactions, incubated 2 min at room temperature, washed twice with freshly prepared 80 % EtOH using a magnetic rack, and eluted with water.

DNA was quantified by Nanodrop, diluted to 5 ng/μL, quantified by Qubit, diluted to 0.4 ng/μL, quantified by Qubit and diluted to 0.2 ng/μL for library preparation. Library preparation was done with the Nextera XT DNA kit (Illumina, cat. FC-131-1096) which fragments input DNA and adds sample-specific barcodes by tagmentation. Although we used one barcode per sample, it is also possible to pool amplicons before library preparation and use the same barcode for multiple samples provided that samples don’t need to be identified individually or that reads for each sample can be distinguished after mapping (e.g. non-overlapping amplicons from different genes). Libraries were analyzed with a Tapestation D1000 ScreenTape system (Agilent) or Bioanalyzer HS DNA kit (Agilent), and showed an average fragment size of around 500 bp (range 400 – 600 bp). Average fragment size, together with the DNA concentration measured with Qubit, was used to determine molarity and an equimolar pool of libraries was prepared. This pool was again analyzed using Tapestation or Bioanalyzer, measured by Qubit and diluted to 2 nM as input for the Illumina sequencing workflow. The library pool was then sequenced using 150 bp reads with a Miniseq Mid Output kit, 2×150 cycles (Illumina, cat. FC-420-1004), or a Nextseq 500 V2 Mid Output kit, 150 cycles (Illumina, cat. FC-404-1001).

### Targeted mRNA sequencing (*lin-41*)

Mutated F2, arrested at the L 1 developmental stage, were obtained from Cas9-induced P0 as described above. 40,000 were directly frozen for genomic DNA extraction. 80,000 were directly frozen for RNA extraction by adding 1 mL TRIzol reagent (Thermo Fisher Scientific, cat. 15596-018), homogenization with a Precellys 24 tissue homogenizer (Bertin Instruments) and storage at -80°C. 5,000 L1s were seeded on large 15 cm NGM plates at 24°C and collected 32 hours later, at late-L4 stage, and prepared for RNA extraction like the L1 sample. At 32 hours, lin-41 mRNA is fully downregulated^35^, while the lethal vulva bursting occurs later after molting, in the adult stage^24^.

RNA was chlorophorm-extracted as follows. Samples were thawed, 0.2 mL of chlorophorm added, incubated for 3 minutes, and centrifuged for 15 minutes at 12,000 x g at 4°C. The upper aqueous phase was transferred to a new tube, 2 μL GlycoBlue (30 μg) were added, 500 μL of isopropanol were added and sample was incubated for 10 minutes. Sample was centrifuged 10 minutes at 12,000 x g at 4°C, supernatant discarded, and 1 mL of 75% EtOH was added. Sample was centrifuged for 5 minutes at 7,500 x g at 4°C, supernatant removed, pellet air-dried and resuspended in 20 μL RNase-free water. RNA concentrations ranged between 1,000 - 2,000 ng/μL, as determined on a Nanodrop ND-1000. Sample was diluted to 300 ng/μL and used for reverse transcription.

RNA was reverse transcribed using Maxima H Minus Reverse Transcriptase (Thermo Fisher Scientific, cat. EP0752). A reaction containing 11.5 μL RNA (3.45 μg), 2 μL gene-specific RT primer at 10 μM (oJJF890 “3’end”, containing a UMI and PCR handle), 1 μL dNTP Mix (10 mM each), was incubated 5 minutes at 65°C. Then 4 μL 5X RT buffer, 0.5 μL RiboLock RNase inhibitor, and 1 μL (200 U) Maxima H Minus reverse transcriptase were added and the reaction was incubated for 30 minutes at 60°C, and 5 minutes at 85°C.

PCR was performed with a *lin-41*-specific primer containing a sample-specific barcode (oJJF1140-1147 for samples N2, 1516, 2627, pool3 at L1 and L4 stages) binding in the second last exon and a primer (oJJF960) binding the PCR handle introduced by the reverse transcription primer. 2 μL of each RT reaction was used as template in 4 PCR reactions, each containing 10 μL 5X HF buffer, 1 μL dNTP mix (10 mM each), 5 μL F+R primer mix (10 μM), 0.2 μL Phusion polymerase, 32 μL water and 2.5 μL DMSO (5% final). Samples were incubated at 98°C 3 min, followed by 35 cycles of 98°C 10 sec, 69 °C 20 sec, 72 °C for 1 min with a final elongation at 72 °C for 7 min. PCR was then analyzed on an agarose gel and DNA was cleaned up using Ampure XP beads (Beckman Coulter, cat. A63881). For this the four PCR reactions were pooled resulting in 100 μL. 80 μL beads were added, incubated for 5 min at room temperature, washed once with 70% ethanol, and DNA was eluted in 10 μL water. This resulted in concentrations between 40-110 ng/μL. All samples were diluted to 40ng/ μL and then pooled. 32 μL of this pool (1280 ng) was then used as the input for SMRTBell (Pacbio) library preparation according to the instruction manual and sequenced using a Pacbio Sequel II sequencer.

### DNA sampling over generations (*lin-41*)

Mutated F1 samples were obtained as described above using large-scale mutagenesis by Cas9 heat shock induction. For this we used N2 as control and 3 lines with sgRNAs against the lin-41 3’ UTR (sg15+sg16, sg26+sg27, sgPool). We conducted the experiment at 16°C and 24°C. 3000 L1 stage animals (F1 generation) were seeded on medium plates with OP50. After egg laying and hatching of the next generation (F2) after 3 or 5 days (24°C or 16°C) F1 and F2 were separated. For this, animals were washed from plates in a final volume of 2 ml M9 buffer into 2 ml Eppendorf tubes. Adult animals sink faster and after circa 2-5 minutes are collected at the bottom of the tube, while L1 animals still swim. This was carefully monitored visually. When most adults (95%) had sunken to the bottom, supernatant M9, containing L1 stage animals, was removed to a separate tube. This was repeated three times by adding 2 ml M9 and separation by sinking. Adult animals were frozen for genomic DNA extraction in circa 20 uL M9. For generations F2-F4, 2000 L1 were seeded on new medium plates, and frozen as adults after separation from the next generation. Generation F5 was frozen at L1 stage. Genomic DNA extraction and targeted large amplicon sequencing was performed as described above.

### Screen for regulatory sequences affecting phenotype

We targeted 8 genes with known RNAi-phenotypes (*dpy-2, dpy-10, egl-30, rol-6, sqt-2, sqt-3, unc-26, unc-54*) using different sets of sgRNAs against regulatory regions. We used lines in which we targeted the 3′ UTR and for some genes we used additional lines targeting predicted enhancer, TATA-box, initiator (INR) and upstream/promoter regions. A list with all samples can be found in a supplementary table.

For each transgenic line we screened 35,000 - 45,000 F2 animals produced from P0 with transiently induced Cas9 expression as described above. Animals were seeded onto NGM plates with food at a concentration of 15,000 per 15 cm plates or at 2,500 - 5,000 per 10 cm plates. Plates were kept at 16°C or 24°C. We then directly screened these plates by eye. Additionally, we collected worms in M9 and dispensed worms in drops on an empty plate. We then observed worms moving in M9 and moving away after M9 was dried (<1 min.). Dpy, Unc, and Rol worms were identified by morphology, their movement in M9 or slow and otherwise impaired movement away from the spot of dispension. Potential mutants were then picked and kept on plates for 2 - 4 generations at 24°C to achieve homozygosity. Animals were then singled by phenotype and genotyped. This resulted in isolation of several mutant strains with the same genotype. We could not distinguish between cousins/siblings coming from the same F1/F2 or independent mutants coming from independently mutated F1s. In these cases, we kept one representative strain. We determined that penetrance was complete for all alleles except for the *sqt-2* locus (n>300 animals). For sqt-2 the penetrance varied between 10-100%. We scored the expressivity of the phenotypes into three categories (+, ++, +++) (n>300 animals). All the reported phenotypes have been determined and validated for several generations at 24°C. We also validated the absence of the extra-chromosomal transgenes judged by the red fluorescent co-injection marker. For *sqt-3* all isolated Dpy animals, characteristic for complete loss-of-function, contained large mutations affecting the coding frame. We therefore screened mainly for reduction-of-function alleles by screening for Rol animals. Non-Rol revertants of the *sqt-3(ins)* Rol animals were isolated using the small-scale approach on 6 cm plates with injection mixes imJJF215 or imJJF230.

### Genotyping

Single worms were picked using a platin wire picking tool and immersed in 10 μL of worm lysis buffer (WLB) (10mM Tris pH 8.3, 2.5 mM MgCl_2_, 50mM KCl, 0.45% NP-40, 0.45% Tween-20, 0.01% gelatine, and freshly added 100 μg/mL proteinase K). Samples were frozen at -80°C for at least 10 minutes, incubated at 60°C for 30-60 minutes, and 95°C for 15-30 minutes in a thermocycler. 1 μL of lysate was used as template in the following PCR. 25 μL PCR reactions were set up as follows. Phusion HF polymerase (New England Biolabs, cat. M0530L) 0.1 μL, 5X HF buffer 5 μL, dNTP mix 0.5 μL, forward and reverse oligos at 10 μM 2.5 μL, water 16 μL, and template DNA. 98°C 3 min, followed by 35 cycles of 98°C 15 sec, 58-72 °C 30 sec, 72 °C for 7 min with a final 7 min at 72 °C. 2 μL of the reaction was then analyzed on an agarose gel. DNA was then cleaned up using AMPure XP Reagent (Beckman Coulter, cat. A63881) by adding 0.8 x volume of beads to 23 μL PCR reaction, 2 min at room temperature, washed twice with freshly prepared 80 % EtOH using a magnetic rack, and eluted with 18 μL water. DNA was then either analyzed by T7 nuclease assay or directly sent to Sanger sequencing. T7 nuclease assay was performed on cleaned up DNA using T7 endonuclease.

### mRNA quantifications with Nanostring and RT-qPCR (*sqt-3*)

10 k L1-arrested synchronized animals were dispensed on 10 cm NGM plates with *Escherichia coli* OP50 at 24 °C. Worms were then collected at different time points (22, 24, 26, 28, 30, 32 hrs), washed once with M9 and homogenized in 1 mL of TRIzol reagent (Thermo Fisher Scientific, cat. 15596-018) using a Precellys 24 tissue homogenizer (Bertin Instruments). RNA was isolated by standard phenol-chlorophorm extraction. RNA expression was quantified using an nCounter (Nanostring) which measures absolute RNA amounts using a set of gene-specific probes. Raw counts were normalized using reference genes (“house-keeping”). For RT-qPCR of pre-mRNA and mRNA we used RNA from the 26 hrs time point where *sqt-3* expression peaked. Pre-mRNA was specifically detected using intron-overlapping primers, while mRNA primers overlapped with exon-exon junctions. Controls without reverse transcriptase (“RT-”) were done to ensure specific amplification of cDNA and no amplification from potential contaminating genomic DNA. Final values were obtained by normalizing to pre-mRNA or mRNA of *tbb-2* and presented relative to N2 wild-type controls. Probes and primers can be found in a supplementary table.

### Transplantation of insertion sequence into the *dpy-10* and *unc-22* 3′ UTRs

Knock-in animals were produced using Cas9/tracRNA/crRNA RNP injections with ssDNA oligo repair templates. Injection mixes contained: 0.3 ug/ul Cas9 protein (Alt-R Cas9 V3 from IDT, cat. no 1081058), 0.12 M KCl, 8 nM Hepes pH 7.4, 8 uM tracrRNA (Alt-R from IDT, cat no 1072532), 8 uM crRNA (custom crRNA, Alt-R from IDT), 3.15 ng/ul pJJF062 (GFP co-injection marker), 3.15 ng/ul pIR98 (HygroR), 0.75 uM of a ssDNA oligo repair template, in duplex buffer (IDT). To prepare injection mixes, Cas9 protein was mixed with KCl and Hepes. crRNA and tracrRNA were annealed in duplex buffer by 5 min at 95 °C and ramp down to 25°C and added. Cas9/tracRNA/crRNA mix was incubated at 37°C for 10 min. Then plasmids and ssDNA repair template were added and 10 P0 animals were injected. For each injection mix 8 F1s positive for the co-injection marker were picked and genotyped using two PCR reactions (one primer pair flanking the insertion, the other with one primer binding in the insertion).

### CRISPR-Downstream Analysis and Reporting Tool (“crispr-DART”)

The source code along with installation instructions and sample input files can be accessed here: https://github.com/BIMSBbioinfo/crispr_DART. The source code for reproducing the figures generated based on the pipeline’s output can be found here: https://github.com/BIMSBbioinfo/froehlich_uyar_et_al_2020.

### Purpose of the crispr-DART pipeline

In order to evaluate the outcomes of the CRISPR-Cas9 induced mutations by the protocol described in this study, we developed a computational pipeline to process the high-throughput sequencing reads coming from samples treated/untreated with CRISPR-Cas9. Although we developed the pipeline to address the hypotheses considering the specific experimental design in this study, we tried to make the pipeline as generic as possible to accommodate different experimental setups, hoping that the pipeline can be useful to the scientific community carrying out genome editing experiments using the CRISPR-based technologies, in particular those that aim to introduce many combinations of mutations in a genome via inducing double-stranded DNA breaks repaired via the non-homologous end joining pathways. The pipeline can handle both short (single- or paired-end Illumina) and long reads (PacBio) from both DNA and RNA, so that the effects in the DNA editing can be also observed in the matching RNA samples. Each sample can contain multiple sgRNAs targeting multiple regions of the genome. A sample might contain reads coming from one or more individual genomes/transcriptomes. However, each sample is treated as if it is a single individual with a mosaic genome. The first purpose of the pipeline is to serve as a quality control/reporting tool to evaluate the genome-editing experiment and address the following questions: Has the CRISPR-Cas9 treatment induced any mutations? If so, how are they distributed in the genome? Do the mutations that are commonly found in many reads originate at the intended cut site based on the designed guide RNA matching sites in the genome? How efficient were different guide designs in inducing DNA damage? Can we capture long deletions if there are multiple sgRNAs used in the same sample targeting nearby sites? How diverse are the deletions or insertions detected at the cut sites? We developed the pipeline to produce HTML reports collated into a website with interactive figures that help the user to quickly visualize and evaluate the outcomes of their experiment.

The second purpose of the pipeline is to produce many processed files containing information that can be useful for further analysis by external tools. Therefore, the pipeline’s output consists of BAM files, bigwig files, BED files, and many different tables containing information about insertions and deletions along with the reads in which they were detected. In this study, many of the figures made for the manuscript were generated based on these intermediate files to address the many custom questions.

### Input description

crispr-DART is implemented using Snakemake^95^ following the practices as implemented for the PiGx pipelines^96^. The input consists of a settings file in yaml format, which contains configurations for the tools used in the pipeline. Moreover, it contains file paths for where the sequencing reads are located, the target genome sequence to be used for mapping the reads, the sample sheet file (in comma-separated file format) which contains the experimental design, the file containing the genomic coordinates of the expected sgRNA cut sites (in BED file format), and a table (in tab-separated format) that is needed for when a pair of samples are to be compared (for instance to observe the differences in per-base distribution of deletions detected in a treated sample and an untreated control sample).

### Steps of the pipeline

The pipeline consists of the following sequence of processing steps (see **Supplementary Figure 2b**):

### Pre-processing reads

- Quality control (using fastqc^97^ and multiqc^98^ and quality improvement of reads (using Trim-Galore!^99^)

### Mapping

- Mapping/alignment of the reads to the genome (using BBMAP^100^). We use BBMAP for read alignment because it can handle both long and short reads, both single-end and paired-end reads, both DNA and RNA reads, and it can help detect both short and long insertion and deletion events.
- Re-alignment of reads with indels using GATK^101^. This step helps reconcile different indel alignments to minimise the noise in alignments.

### Finding indels and generating indel related statistics

- Extraction of indels from the BAM files using R packages GenomicAlignments^102^ and RSamtools^103^, producing:
  - BED files for the genomic coordinates of insertions and deletions
  - Bigwig files for :
    □ Alignment coverage to see how many reads
    □ Insertion/deletion/indel aligned per each base of the genome. scores: this represents the ratio of reads with an insertions, deletion or either (indel) to the number of reads aligned at a given base position of the genome. These files are very useful in visualising profiles of the degree of the mutation damage at per-base resolution.

- Tab-separated format files for:
  □ Inserted sequences: this table contains the list of all reads with an insertion, along with the exact genomic coordinate of where the insertion occurs, and the actual sequence of the inserted segment.
  □ Indels: This table contains the genomic coordinates of the deletions and insertions along with how many reads actually support the insertions/deletions and the maximum depth of coverage (considering all reads) along the deleted segment or at the insertion site. This table contains all insertions/deletions supported by one or more reads. list of reads with insertions/deletions along with the coordinates of the insertions/
  □ Reads with indels: This table is the complete deletions.
  □ sgRNA efficiency: This table contains statistics about the efficiency of each guide RNA in inducing mutations at the targeted site of the genome. The efficiency of a sgRNA is defined as the ratio of the number of reads with an insertion/deletion that start or end at +/- 5bp of the intended cut-site to the total number of reads aligned at this region.

### Generating a website of HTML reports

All the pre-processed files from the previous steps are combined to generate interactive (where applicable) HTML reports from all the analysed samples that exist in the input sample sheet. For each targeted region (assuming a region of a few thousand base pairs that is sequenced), currently four different reporting Rmarkdown scripts are run. The resulting HTML files are organised into a website using the ‘render_site’ function of the Rmarkdown package^104^. Thus, all the processed data and outcomes can be quickly browsed through a website. The resulting website that is the output of the pipeline for this study can be browsed here: https://bimsbstatic.mdc-berlin.de/akalin/buyar/froehlich_uyar_et_al_2020/reports/index.html

### How to reproduce the figures in the manuscript

The source code and usage instructions for reproducing the figures in this manuscript, that were made based on the output of the crispr-DART pipeline, can be found here: https://github.com/BIMSBbioinfo/froehlich_uyar_et_al_2020.

### RNA analysis: Comparing deletions in L1 and L4 stages

Deletions in PacBio L1 and L4 samples were filtered to keep only those deletions supported by at least 5 PacBio reads. Each deletion was categorized based on their overlap with important sites in the 3’UTR of lin-41:

- First let-7 micro-rna binding site (only the seed region) (LCS1_ seed): chrI:9335255-9335263
- Second let-7 micro-rna binding site (only the seed region) (LCS2_ seed): chrI:9335208-9335214
- First let-7 micro-rna binding site minus the seed region (LCS1_3compl): chrI:9335264-9335276
- Second let-7 micro-rna binding site minus the seed region (LCS2_3compl): chrI:9335215-9335227

Deletions were further categorized based on whether they overlap both let-7 micro-rna seed regions, and also those that don’t overlap any of these defined regions.

Deletion frequency values were computed and the ratio of deletion frequencies between L4 stage and L1 stage samples were computed in log2 scale. For each category of deletions, a wilcoxon rank-sum test was computed to test the null hypothesis that the stage specific abundance of deletions that overlap a let-7 binding site is not different from those deletions that don’t overlap any of these sites.

### RNA analysis: Clustering pacbio reads and comparing genotype abundance in L1 and L4 stages

Reads from both L1 and L4 stage lin-41 RNA samples sequenced using PacBio that cover the region between chrI:9334840-9336100 (the region from the beginning of the amplified segment up to the first intron) were selected to make sure that all reads that go into analysis are covering the whole segment. For each read, the alignment of the read (including the inserted sequences) were obtained and all combinations of k-mers (k=5)were counted within these alignments allowing for up to 1 mismatch using Biostrings package^105^. Seurat package^106^ was used to process the k-mer count matrix to do scaling, dimension reduction (PCA and UMAP) and network-based spectral clustering. The clustering of long PacBio reads covering the region enabled us to cluster reads into genotypes, thus taking advantage of the length of the reads while also allowing for the high rate of indels in the PacBio reads (compared to the Illumina reads).

### Fitness analysis

For this analysis, we have utilized lin-41 DNA samples sequenced (Illumina single-end sequencing) from multiple generations from F2 to F5 of the same pool of animals treated with sgRNA guides “sg15 and sg16”, “sg26 and sg27” and “sg pool”.

### The important sites considered for this analysis are

- First let-7 micro-rna binding site (only the seed region) (LCS1_ seed): chrI:9335255-9335263
- Second let-7 micro-rna binding site (only the seed region) (LCS2_ seed): chrI:9335208-9335214
- First let-7 micro-rna binding site minus the seed region (LCS1_3compl): chrI:9335264-9335276
- Second let-7 micro-rna binding site minus the seed region (LCS2_3compl): chrI:9335215-9335227
- Poly-adenylation signal: chrI:9334816-9334821
- Stop-codon: chrI:9335965-9335967

We would like to address the question whether the deletions that exist at F2 are exposed to purified selection over generations if they overlap the important sites on the 3’UTR region of lin-41. We do this analysis in two ways:

- First, we count the deletions categorised by their overlap (or non-overlap) with the important sites that exist in F2 generation and see how many of them still exist in later generations.
- Secondly, we do the same analysis at the level of reads: counting the reads with deletions that overlap or not overlap the important sites from generations F2 to F5. When comparing the number of reads, the read counts are normalized by the library sizes (total number of reads in the sample).

### sgRNA efficiency comparisons

For comparing observed efficiencies to published prediction scores and other sgRNA characteristics, these were manually extracted from the CRISPOR web application (http://crispor.tefor.net/)^107^.

### Browser shots

Browser shots were compiled using indel profiles and top indels provided by the computational pipeline as BigWig and BED files and loading them into the UCSC genome browser^108^ or the IGV browser^109^ followed by export as vector graphics compatible format. We used *C. elegans* genome version ce11/WBcel235 including 26 species base-wise conservation (PhyloP).

### Graphics

Final figures were assembled using Adobe Illustrator without changing the plotted data, with adjustments to composition, axis labels, line widths, colors, etc.

### Manuscript formatting

This manuscript was composed using Adobe InDesign starting with a template from https://github.com/cleterrier/ManuscriptTools. We thank Christophe Leterrier for sharing it.

### Data reporting

No statistical methods were used to predetermine sample size. The experiments were not randomized. The investigators were not blinded to allocation during experiments and outcome assessment.

### Strain and plasmid availability

*C. elegans* mutant strains will be available through the Caenorhabditis Genetics Center (CGC) as indicated in a supplementary table. Plasmids generated for this work for heat shock Cas9 expression (pJJF152) and proof of concept sgRNAs (targeting SECGFP, *dpy-10, sqt-3*) will be available from Addgene as indicated in a supplementary table or for specific sgRNAs upon direct request from the authors.

### Data availability

Raw sequencing data can be found at the following link: https://bimsbstatic.mdc-berlin.de/akalin/buyar/froehlich_uyar_et_al_2020/reads.tgz.

Sample sheet which describes the experimental setup for running crispr-DART can be found here: https://bimsbstatic.mdc-berlin.de/akalin/buyar/froehlich_uyar_et_al_2020/sample_sheet.csv. The output of crispr-DART for this data can be found here: https://bimsbstatic.mdc-berlin.de/akalin/buyar/froehlich_uyar_et_al_2020/crispr_dart_pipeline_output.tgz.

### Code availability

The crispr-DART software pipeline can be found at: https://github.com/BIMSBbioinfo/crispr_DARTR scripts to reproduce figures is available at: https://github.com/BIMSBbioinfo/froehlich_uyar_et_al_2020

## Acknowledgements

We are very grateful to Baris Tursun, Luisa Cochella, David Koppstein, and the Rajewsky, Akalin labs for helpful feedback and discussions and Claudia Quedeau, Daniele Franze, the sequencing facility for Pacbio sequencing and Sergej Herzog for technical assistance. We thank the Caenorhabditis Genetics Center (CGC), Christian Frøkjaer-Jensen, Erik Jorgensen, João Ramalho, Mike Boxem, Daniel Dickinson, Bob Goldstein, and Jason Chin for sharing plasmids and strains. B.U. acknowledges funding by the German Federal Ministry of Education and Research (BMBF) as part of the RNA Bioinformatics Center of the German Network for Bioinformatics Infrastructure (de.NBI) (031 A538C RBC).

## Author contributions

J.J.F. and N.R. developed concepts, methodology, and discussed the data. J.J.F. and B.U. performed validation, formal analysis, curation and visualization of data. B.U. wrote the software with input from J.J.F. and A.A. P.G. contributed to software. J.J.F., M.H. and K.T. performed investigation and experimental work. J.J.F. wrote the original draft. J.J.F., B.U., A.A., N.R. reviewed and edited the manuscript. A.A. and N.R. contributed with resources, supervision, project administration and funding acquisition.

**Supplementary Figure 1.**
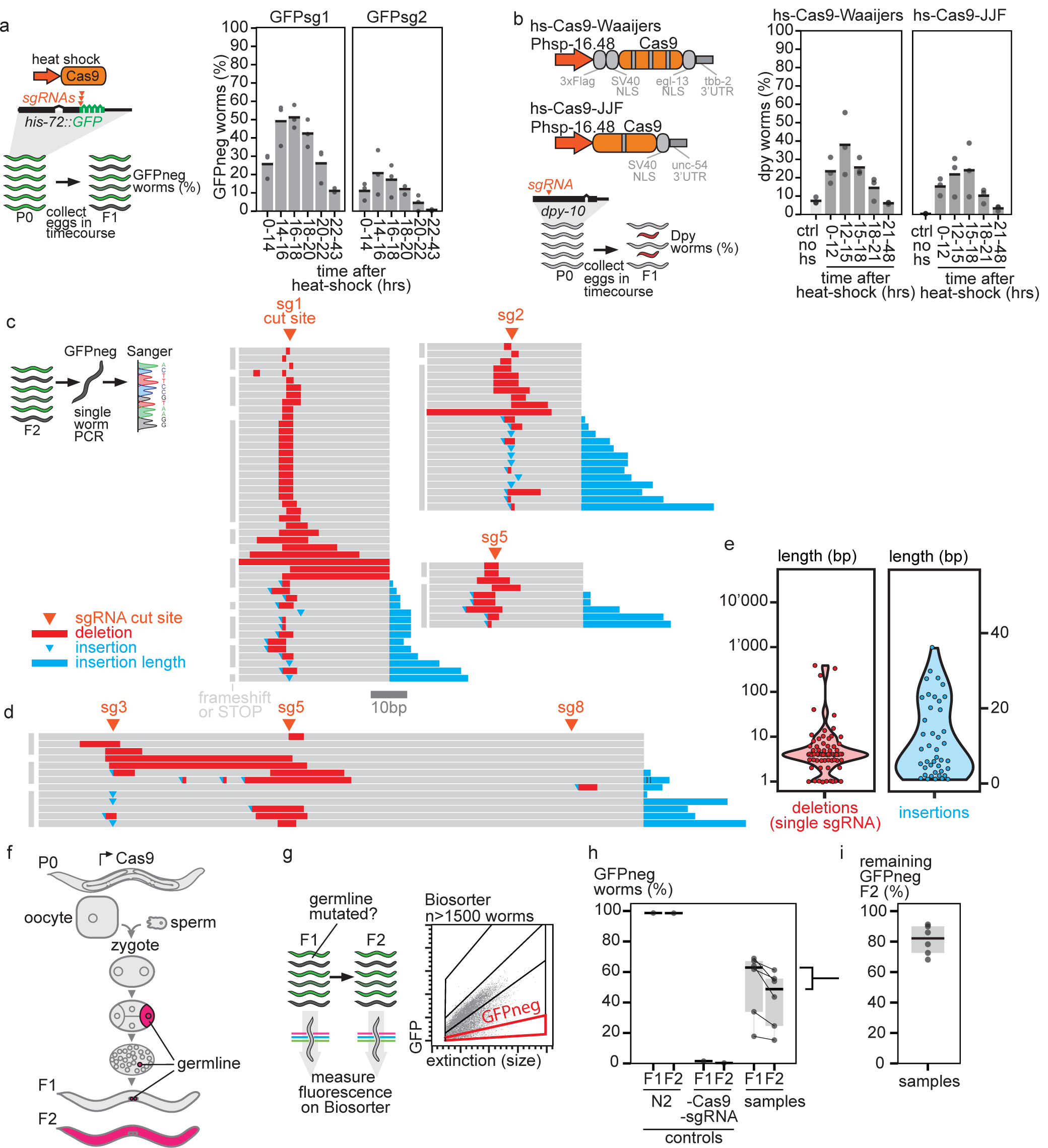
Transiently induced Cas9 expression creates germline indel mutations. **a**, Defining the temporal dynamics of Cas9 induction. An endogenously tagged *his-72::GFP* was targeted with two different sgRNAs. After a two hour heat shock, eggs were collected in a time course and GFP-negative animals were counted. The eggs collected 14 – 16 hrs after heat shock produced the most GFP-negative animals. **b**, Comparison of two different plasmids for heat shock inducible Cas9 in a time course. *Dpy-10* was targeted with a sgRNA, time course was performed as in a) and Dpy progeny were counted. Eggs collected 12 – 14 hrs after heat shock produced the most Dpy animals. Both plasmids performed comparably. **c**, Indel mutations detected by Sanger sequencing of individual GFP-negative animals after targeting *his-72::GFP* with sgRNAs. **d**, Sanger sequencing of indel mutations created by a pool of three sgRNAs. **e**, Length distribution of the indels from individual GFP-negative worms. Deletion length is shown only for the two lines with a single sgRNA. Insertion length is shown for all three lines including the line with a pool of sgRNAs. **f**, A scheme showing the germline lineage in *C. elegans*. F2 animals are created by a germline cell which is determined in the F1 4-cell embryo. **g**, Scheme showing automated fluidics measurement of F1 and F2 GFP negative animals to determine the amount of germline mutations. **h**, Amount of GFP-negative F1 and F2 animals in control strains and after targeting *his-72::GFP* with sg1, sg2, pool1 or pool2. N = 1,662 - 21,983 analyzed worms per sample. **i**, Difference in the amount of GFP-negative animals between F1 and F2 generation. Almost the same amount (80%) of GFP-negative animals in the F2 generations indicates high germline transmission of mutations.

**Supplementary Figure 2.**
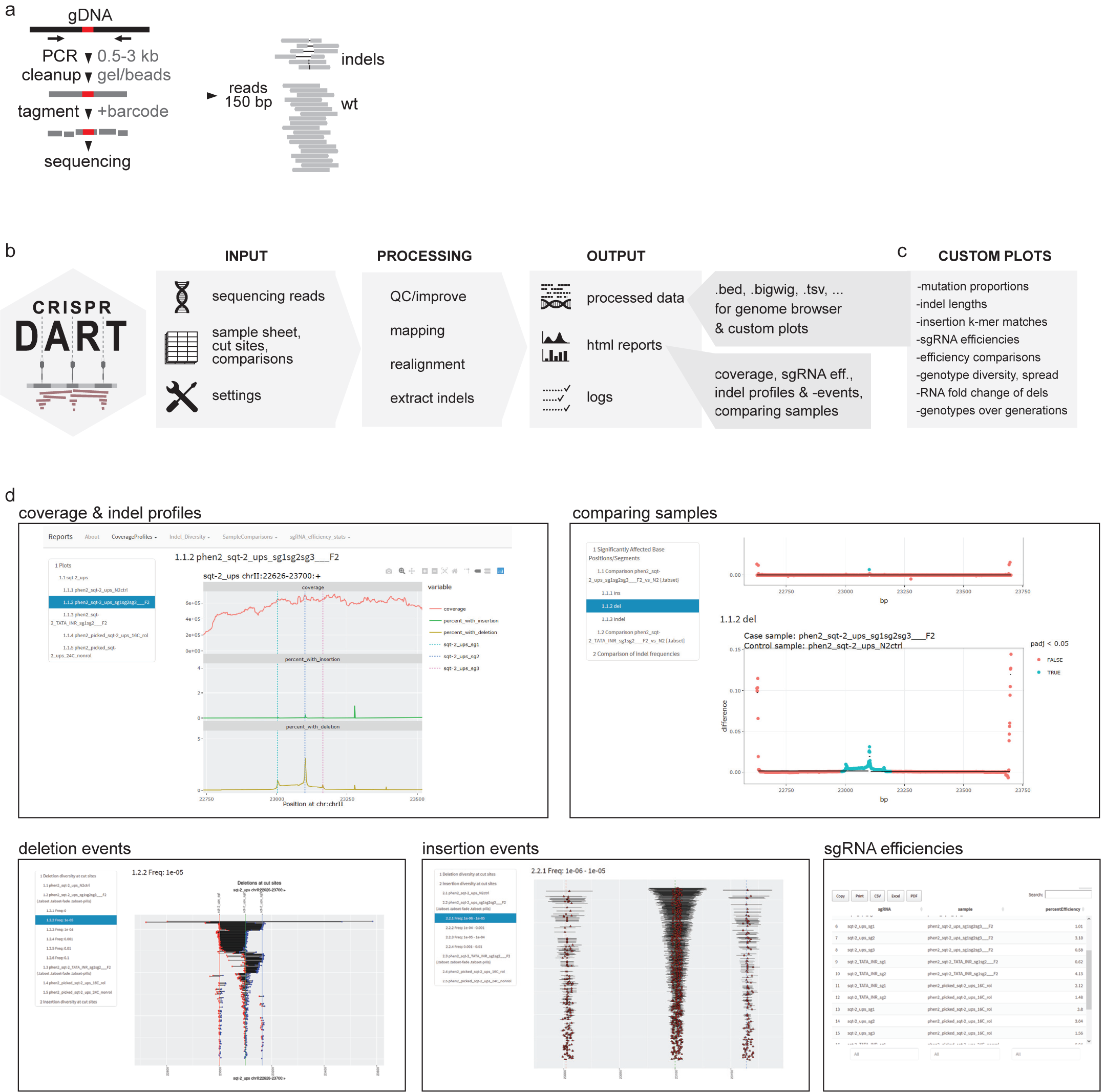
Software pipeline “crispr-DART”. **a**, Scheme showing our long amplicon sequencing approach. **b**, Diagram of the software pipeline “crispr-DART”. The user provides input files and the pipeline produces processed genomic files and html reports. **c**, Custom analyses for this study were then performed with R scripts using the processed genomic files as input. **d**, Screenshots of the html reports provided by crispr-DART. For “crispr-DART” and R scripts see “Code Availability” in the Supplementary Methods.

**Supplementary Figure 3.**
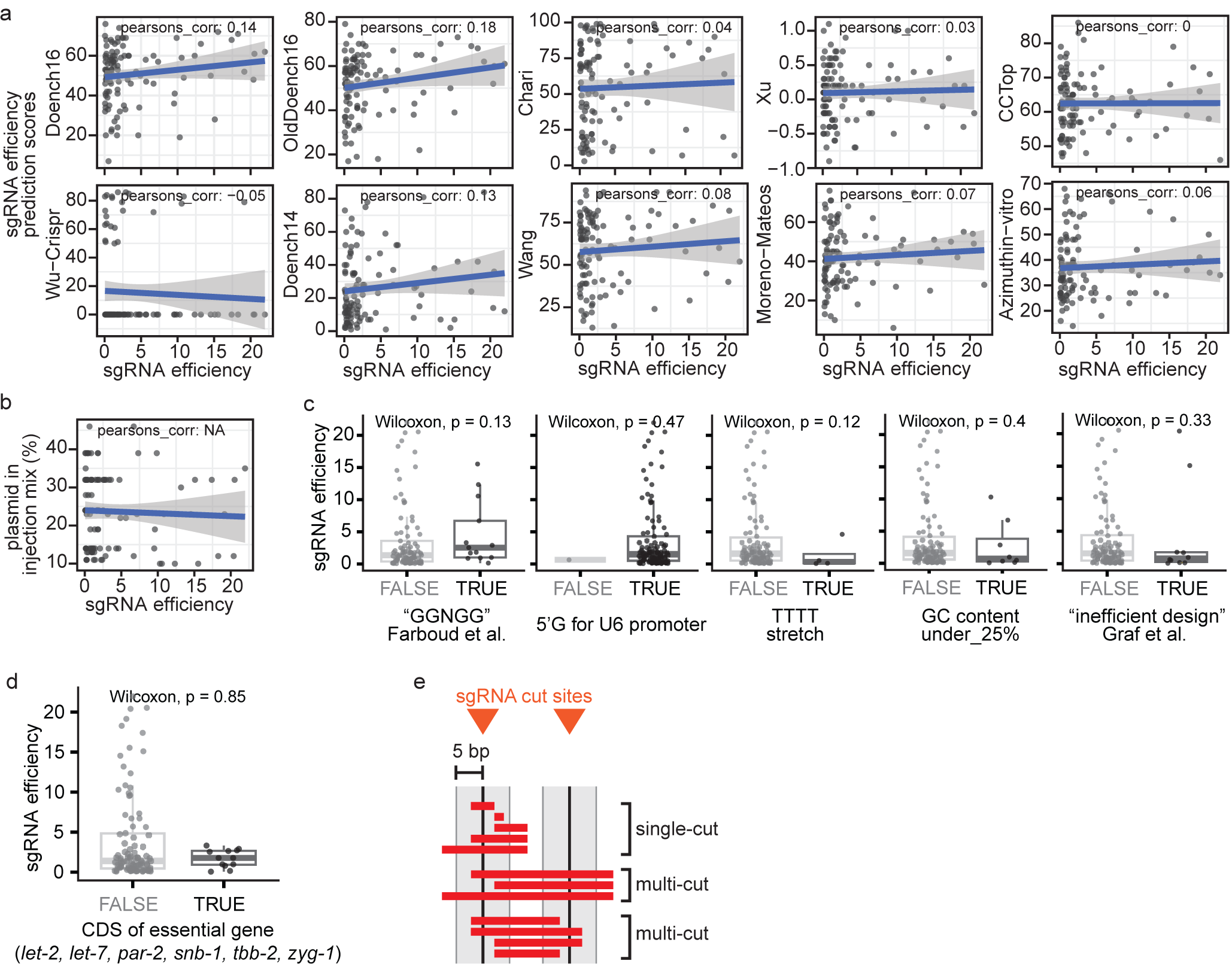
sgRNA efficiency characteristics. **a**, Correlation of various prediction scores for sgRNA efficiency and our observed sgRNA efficiency (n=91 sgRNAs). **b**, Correlation of the percentage of plasmid in the original injection mix and the observed sgRNA efficiency. **c**, Comparison of sgRNA efficiency for different sgRNA features. **d**, Comparison of sgRNA efficiency for sgRNAs targeting the coding sequence of essential genes and all other sgRNAs. No difference could be observed, indicating that low efficiencies for specific sgRNAs were not due to lethality. **e**, Scheme which illustrates how single-cut or multi-cut deletions were categorized computationally.

**Supplementary Figure 4.**
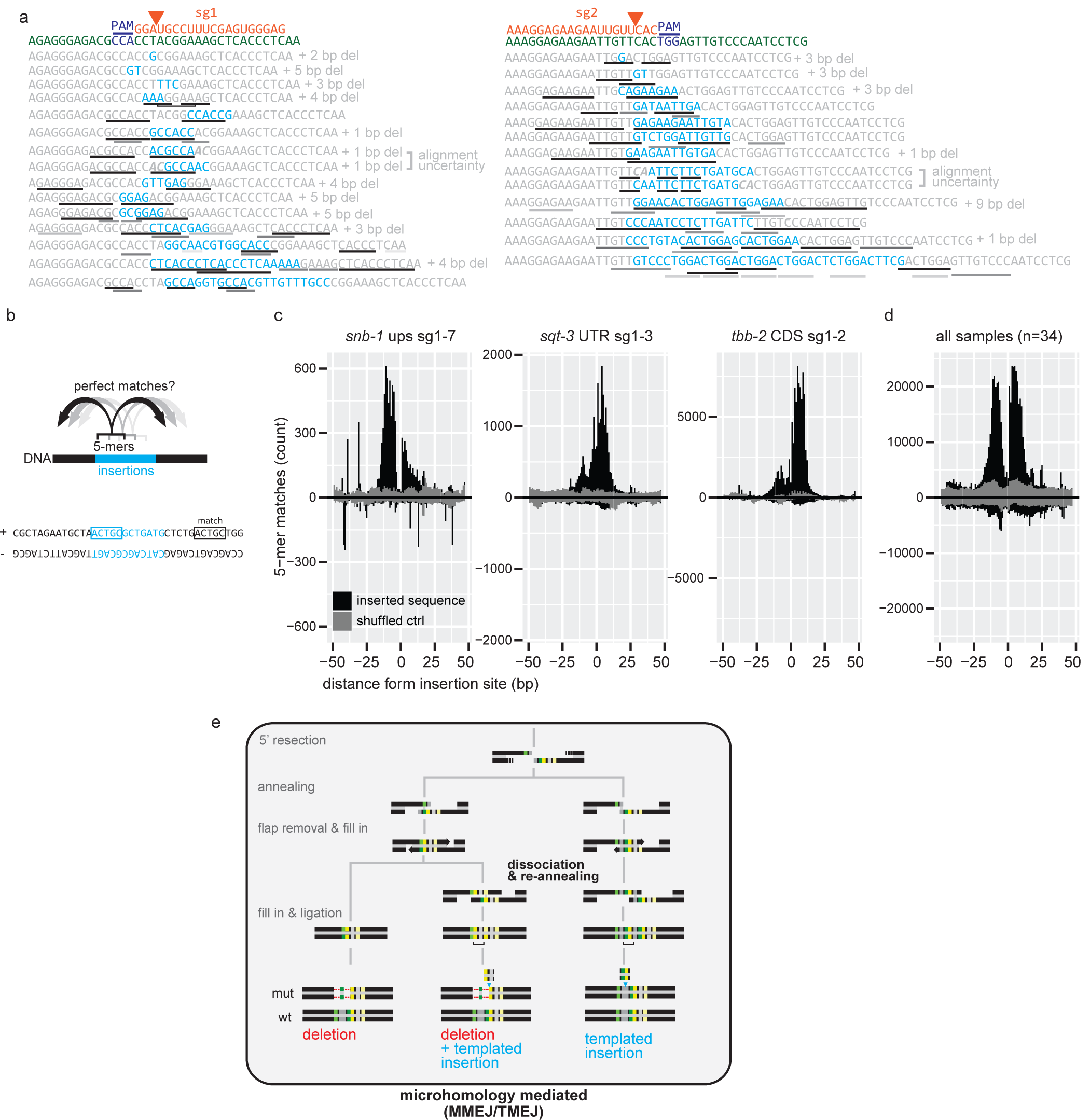
Insertions are templated from surrounding sequences. **a**, Examples of microhomology observed between insertions and surrounding regions in genotypes of GFP-negative *his-72::GFP* animals. **b**, Scheme showing the analysis approach which matches all possible 5-mers from an insertion to the surrounding sequence. **c**, Matches of 5-mers from insertions to surrounding sequence (+/- 50bp), shown for three samples. **d**, For data from 34 samples, matches of 5-mers from insertions to surrounding sequence (+/- 50bp). **e**, Diagram showing mechanistic steps of dsDNA repair via microhomology-mediated end joining. Highlighted is the dissociation & re-annealing step which could lead to the templated insertions observed in our data.

**Supplementary Figure 5.**
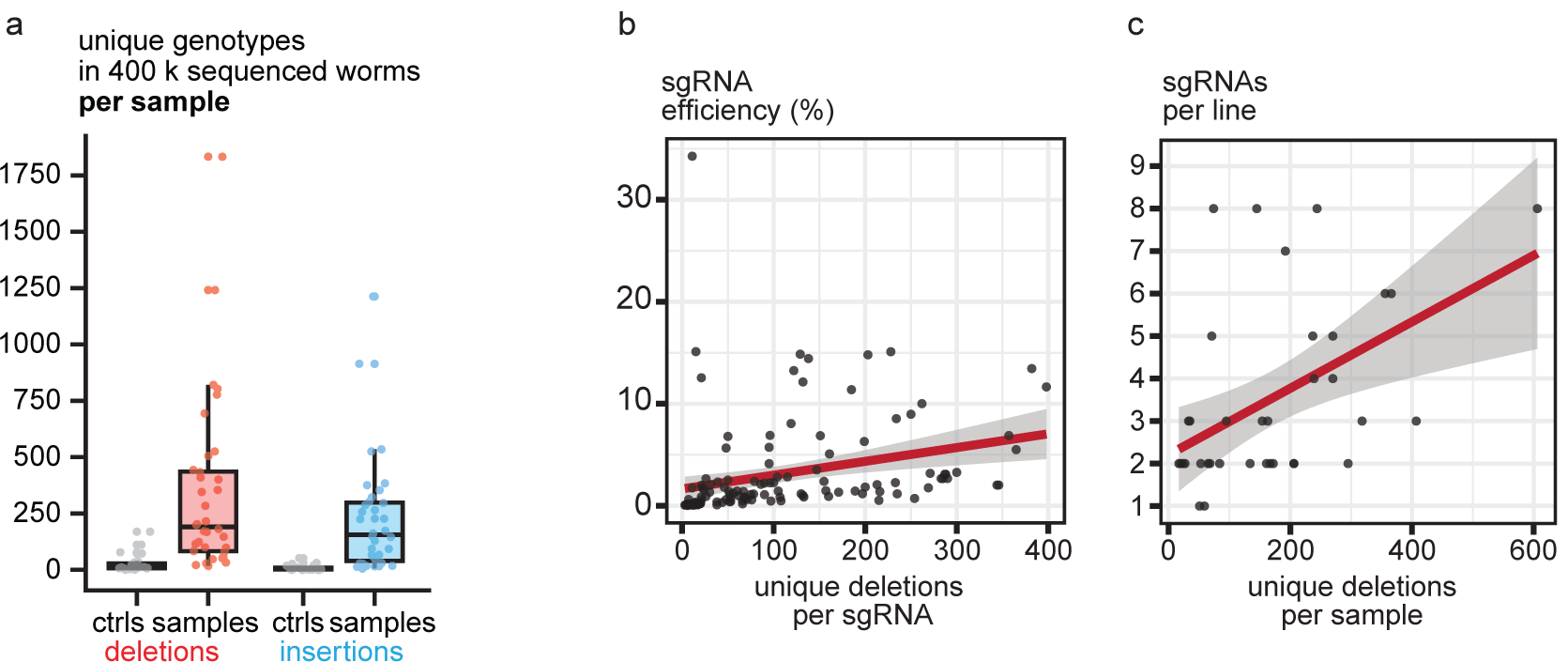
Unique genotypes created by indel mutations correlate with higher sgRNA efficiency and more sgRNAs per line. **a**, Unique genotypes created per sample by deletions or insertions. Each genotype is detected by at least 0.001% of reads mapped to the analyzed position. Ctrls n=24, samples n=36. Wilcoxon, p = 1.7e−08 for deletions, p = 4.7e−09 for insertions. **b**, Correlation between sgRNA efficiency and the created unique deletions per sgRNA (n=91) per sample. **c**, Correlation between the amount of different sgRNAs in a transgenic line and the created unique deletions per sample (n=6084). Unique deletions only from treated samples (n=35), supported by >5 reads and >0.001% frequency.

**Supplementary Figure 6.**
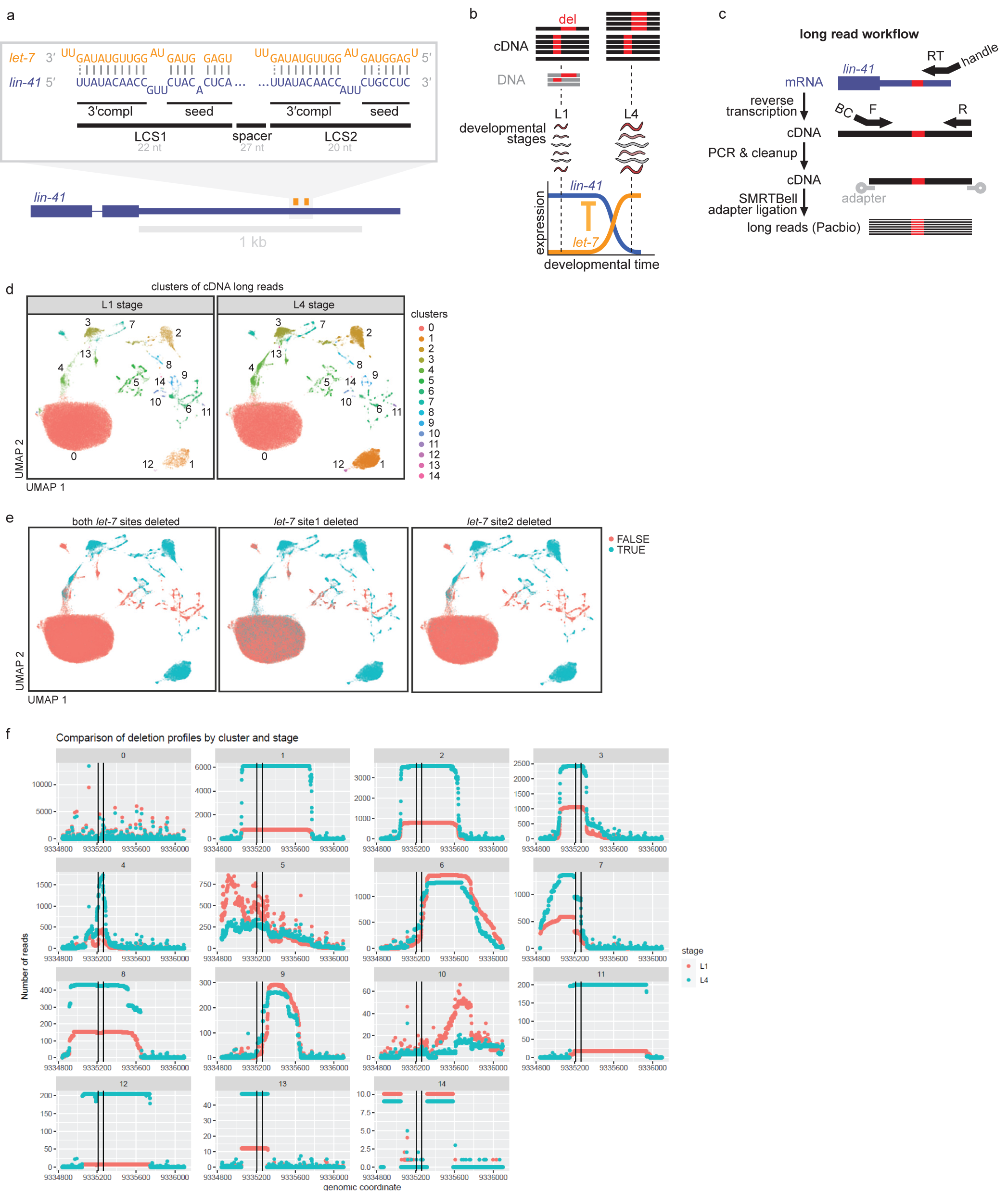
Impact of *lin-41* 3′ UTR deletions on RNA levels. **a**, Diagram showing *let-7* complementary sites LCS1, LCS2 in the *lin-41* 3′ UTR. **b**, Diagram of *lin-41* and *let-7* developmental expression and time points of RNA extraction. **c**, Diagram of the targeted RNA sequencing strategy. cDNA was amplified using a large amplicon and sequenced using the Pacbio long read workflow. **d**, UMAP clusters of long reads covering the complete *lin-41* 3′ UTR, detected in cDNA from L1 or L4 developmental stages. Each dot represents one read. **e**, Status of overlap with *let-7* sites for each read. **f**, Number of detected reads with a deletion (y-axis) per genomic nucleotide (x-axis). Reads are separated by cluster (sub-panels) and developmental stage (L1=red, L4=green). The two vertical black lines indicate the location of the two *let-7* complementary sites (LCS1 and LCS2).

**Supplementary Figure 7.**
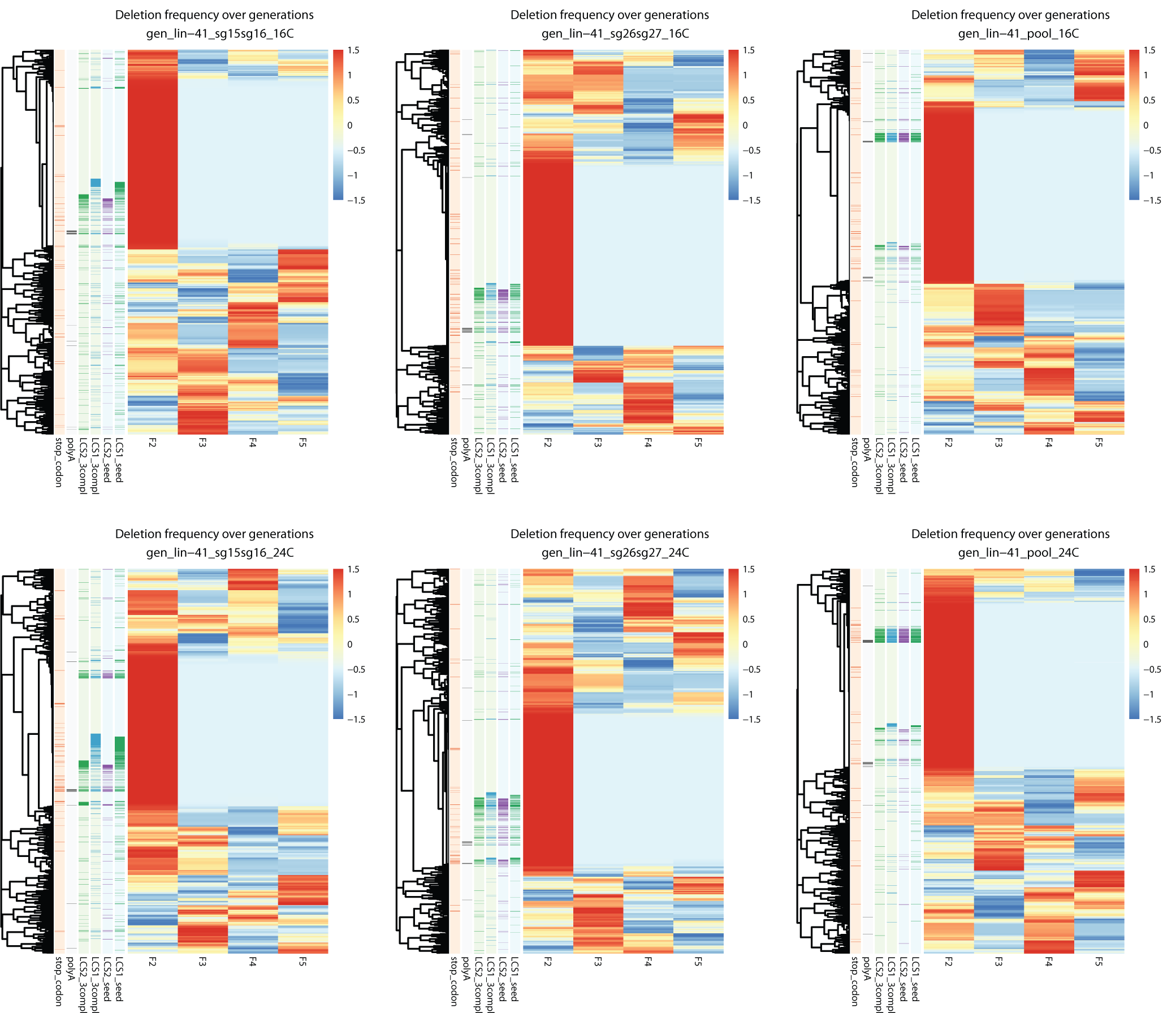
Abundance of *lin-41* 3′ UTR deletions over several generations. **a**, The heatmap (pheatmap package) displays the frequency of deletions (on rows) scaled by row over multiple generations (columns). The annotation columns display which deletions overlap different *let-7* binding sites. Log10_read_count is number of reads in log-10 scale that support the deletions at F2 generation.

**Supplementary Figure 8.**
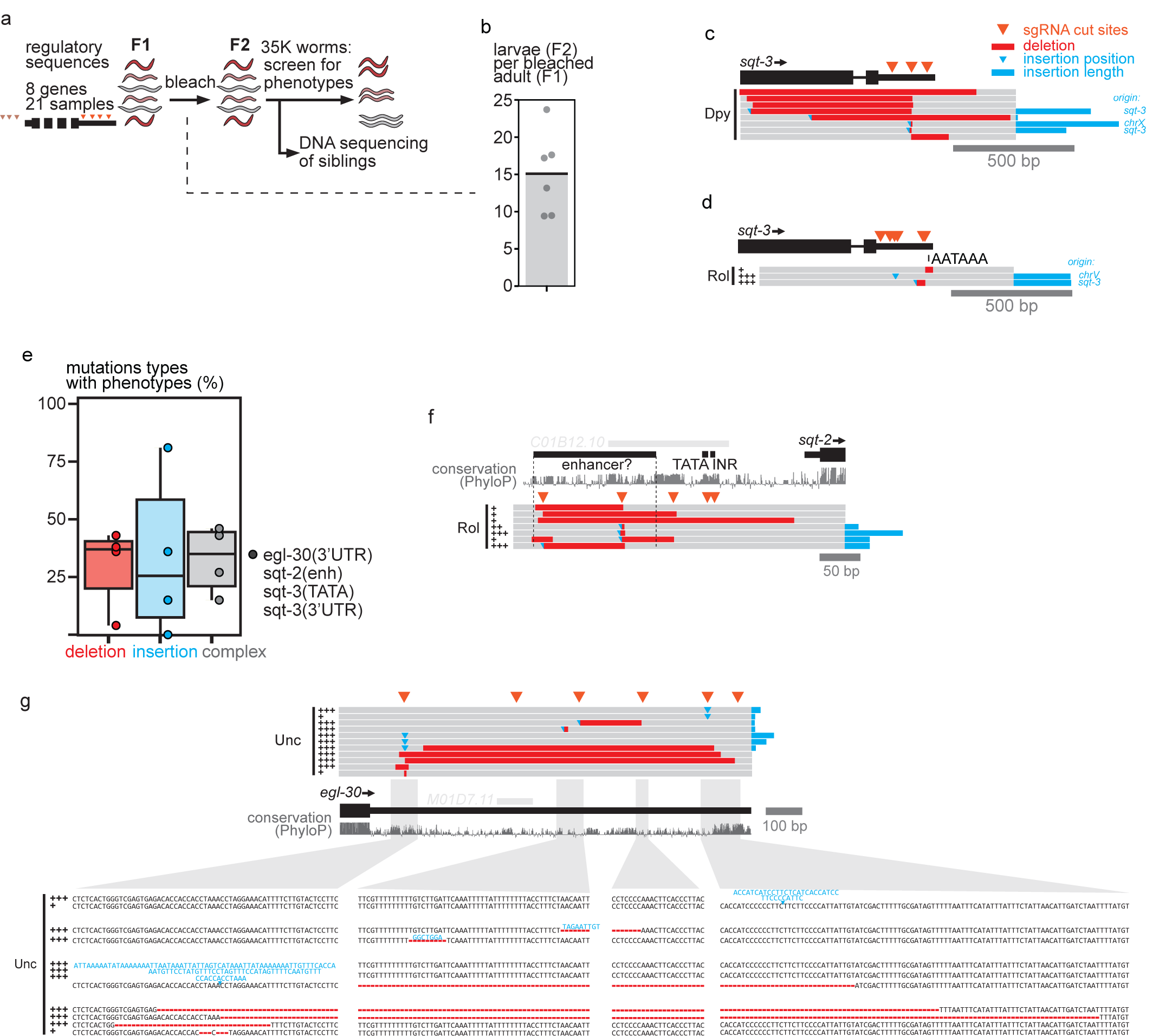
Screen for regulatory mutations with phenotypes. **a**, Outline of the screen. 8 genes were targeted by pools of sgRNAs (between 2-6) in different regulatory regions (some in enhancer, promoter, and all in the 3′ UTR) resulting in 21 samples. For these, 35,000 F2 animals were screened manually for known RNAi phenotypes. The screen allowed us to sequence F2 siblings to determine the mutations present in the screened population. **b**, Amount of F2 progeny obtained per bleached adult of the F1 generation. **c**, Location and extent of mutations affecting the coding sequence in Dpy *sqt-3* mutants. For long insertions the origin was determined by BLAT. **d**, Rol mutations isolated after targeting the *sqt-3* 3′ UTR without sg2. **e**, Mutation types found in reduction-of-function alleles isolated for four targeted regions from three genes (*egl-30, sqt-2, sqt-3*). **f**, Indels affecting a putative enhancer region (Jänes et al. 2018) of *sqt-2*. +, ++, +++ indicate the expressivity of the trait. This was the only region for which penetrance was not complete (10-100%). **g**, Sequences of the mutations in Uncoordinated *egl-30* mutants.

**Supplementary Figure 9.**
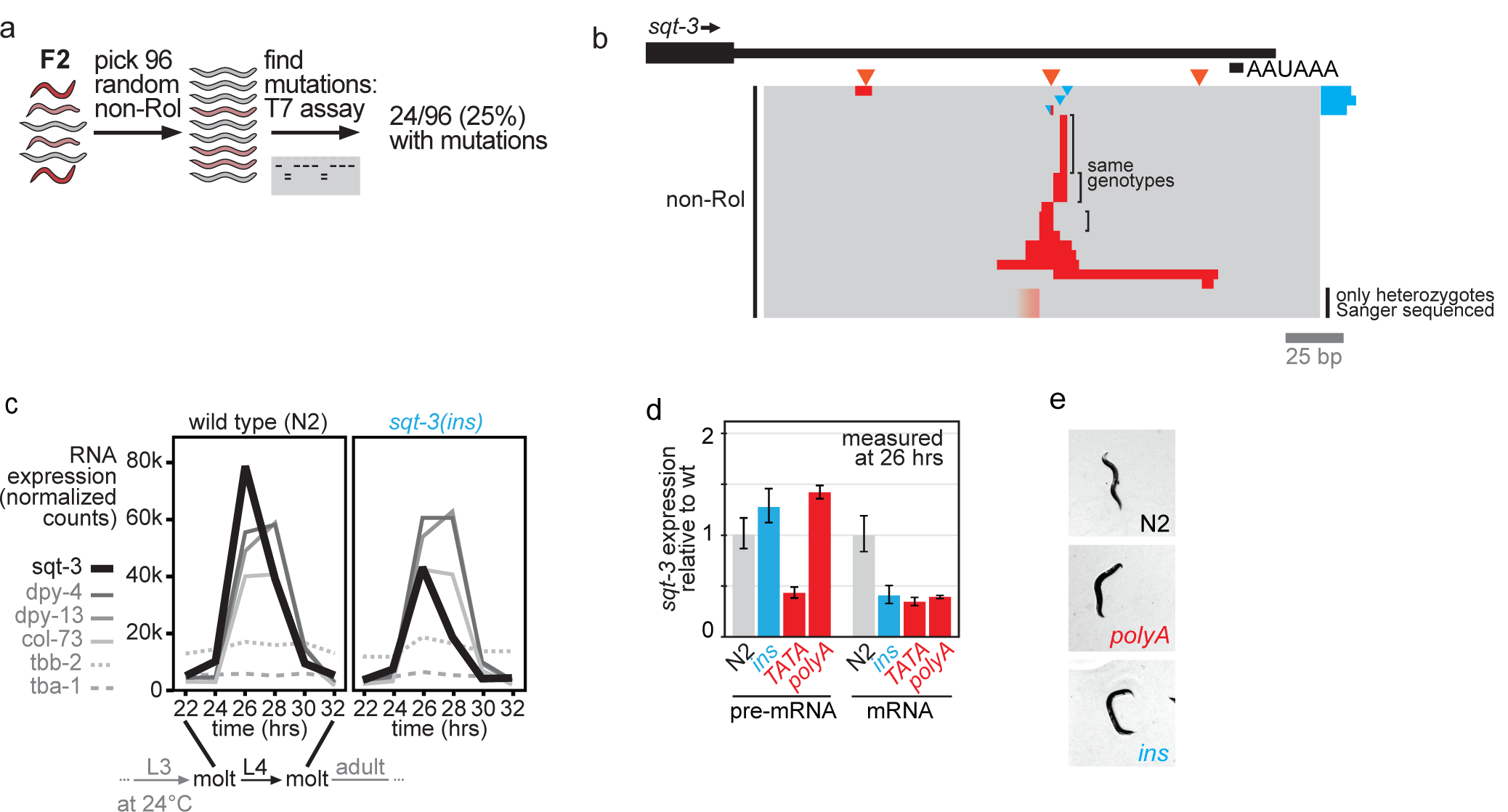
Insertions in the 3′ UTR of *sqt-3* lead to post-transcriptional mRNA reduction. **a**, Isolation strategy of non-Rol mutants. 96 random non-Rol animals were picked and T7 assay was performed to screen for mutations. **b**, Indels in non-Rol animals determined by Sanger sequencing. Exact sequence of two mutations could not be resolved due to heterozygosity. **c**, Quantification of *sqt-3* RNA expression along development during L4 stage in wild type (N2) and mutant. Worms were synchronized by bleaching and RNA was quantified on the Nanostring system. **d**, SQT-3 mRNA or pre-mRNA levels in different *sqt-3* alleles at 26 hrs into synchronized development. Levels were quantified by qRT-PCR with primers specific for the spliced or the un-spliced transcript. **e**, Microscope images of the weak Rol phenotype with only slight bending of the head in the *sqt-3(polyA)* mutant and strong characteristic Rol phenotype in the *sqt-3(ins)* mutant.

**Supplementary Figure 10.**
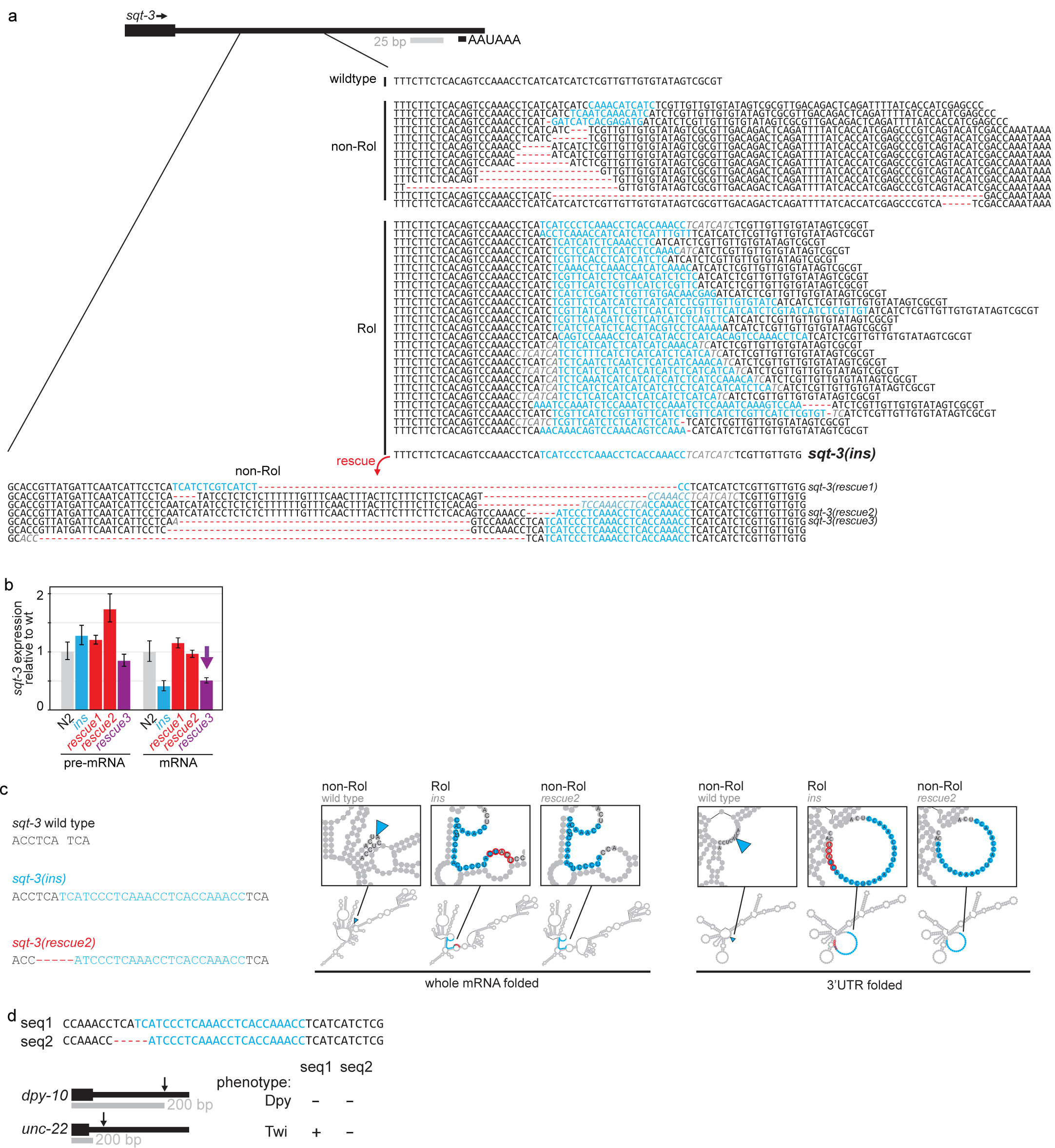
Deletions affecting the repressive RNA element in the *sqt-3(ins)* 3′ UTR can rescue the Rol phenotype. **a**, Nucleotide sequences of relevant 3′ UTR regions in Rol, non-Rol and rescued mutants showing the inserted and deleted nucleotides. **b**, mRNA or pre-mRNA levels of *sqt-3* in mutant and revertants at 26 hrs into synchronized development. Levels were quantified by qRT-PCR with primers specific for spliced or un-spliced transcript. **c**, Predicted RNA secondary structures of wild type, insertion mutant and rescued allele. Predictions were made for the whole mRNA or only the 3′ UTR. **d**, Transplantation of mutant sequences into independent 3′ UTRs. Sequence 1 (from insertion mutant) or sequence 2 (from rescued insertion mutant) were knocked-in at the *dpy-10* or *unc-22* 3′ UTR. Seq1 led to the characteristic reduction-of-function phenotype Twitching (Twi) in *unc-22*.

**Supplementary Table 1:** Overview of samples.

| gene | region | amount of sgRNAs | amplicon size selected |
| --- | --- | --- | --- |
| *lin-41* | CDS | 2 | +/- |
| *lin-41* | 3'UTR | 5 | +/- |
| *lin-41* | 3'UTR | 2 | +/- |
| *lin-41* | 3'UTR | 5 | +/- |
| *lin-41* | 3'UTR | 2 | +/- |
| *lin-41* | 3'UTR | 8 | +/- |
| *lin-41* | 3'UTR | 8 | +/- |
| *lin-41* | 3'UTR | 8 | +/- |
| *lin-41* | downs | 2 | +/- |
| *dpy-2* | 3'UTR | 2 | + |
| *dpy-10* | 3'UTR | 2 | +/- |
| *dpy-10* | 3'UTR | 2 | - |
| *dpy-10* | 3'UTR | 4 | - |
| *egl-30* | 3'UTR | 2 | + |
| *egl-30* | 3'UTR | 2 | + |
| *egl-30* | 3'UTR | 2 | + |
| *rol-6* | enh | 2 | - |
| *rol-6* | TATA | 2 | - |
| *rol-6* | INR | 1 | - |
| *rol-6* | 3'UTR | 2 | - |
| *snb-1* | ups | 7 | + |
| *snb-1* | CDS | 2 | + |
| *snb-1* | CDS | 2 | + |
| *snb-1* | 3'UTR | 8 | + |
| *snb-1* | 3'UTR | 7 | + |
| *sqt-2* | ups | 3 | - |
| *sqt-2* | TATA_INR | 2 | - |
| *sqt-2* | 3'UTR | 3 | - |
| *sqt-3* | TATA | 4 | - |
| *sqt-3* | INR | 2 | - |
| *sqt-3* | 3'UTR | 3 | +/- |
| *sqt-3* | 3'UTR | 3 | - |
| *sqt-3* | 3'UTR | 6 | - |
| *sqt-3* | 3'UTR | 9 | - |
| *unc-26* | 3'UTR | 2 | + |
| *unc-54* | 3'UTR | 3 | + |
| *let-2* | CDS | 2 | - |
| *let-2* | 3'UTR | 6 | - |
| *let-7* | miRNA | 2 | - |
| *par-2* | CDS | 2 | - |
| *tbb-2* | CDS | 2 | - |
| *tbb-2* | 3'UTR | 3 | - |
| *unc-119* | CDS | 1 | +/- |
| *zyg-1* | CDS | 2 | - |
| *zyg-1* | 3'UTR | 3 | - |

**Supplementary Table 2:** Overview of phenotype screen samples.

| gene | dels into coding | isolated mutants | phenotype | region | deletion | complex /insertion | proportion of deletions (%) |
| --- | --- | --- | --- | --- | --- | --- | --- |
| *egl-30* | - | 11 | Unc | 3'UTR | 4 | 7 | 36 |
| *sqt-2* | + | 7 | Rol | enhancer | 3 | 4 | 43 |
| *sqt-3* | + | 13 | Rol | TATA box | 5 | 8 | 38 |
| *sqt-3* | + | 26 | Rol | 3'UTR | 1 | 25 | 4 |
| *dpy-2* | + | 0 | - | 3'UTR | - | - | - |
| *dpy-10* | + | 0 | - | 3'UTR | - | - | - |
| *rol-6* | - | 0 | - | prom, TATA | - | - | - |
| *rol-6* | - | 0 | - | 3'UTR | - | - | - |
| *sqt-2* | + | 0 | - | TATA | - | - | - |
| *sqt-2* | + | 0 | - | 3'UTR | - | - | - |
| *unc-26* | - | 0 | - | 3'UTR | - | - | - |
| *unc-54* | + | 0 | - | 3'UTR | - | - | - |
| *sqt-3(ins)* | + | 15 | Rol->non-Rol | 3'UTR | 11 | 4 | 73 |

